# A global genomic survey of prokaryotic carbon fixation reveals an oxygen-tolerant rTCA cycle in the surface ocean

**DOI:** 10.64898/2026.09.02.748891

**Authors:** Anna Mankowski, Sander Wuyts, Anthony Fullam, David Eckey, Jonas Schiller, Ramiro Logares, Jaime Huerta-Cepas, Thomas S B Schmidt, Michael Kuhn, Peer Bork

**Affiliations:** European Molecular Biology Laboratory, Molecular Systems Biology Unit, 69117 Heidelberg, Germany; ImmuneWatch, 2000 Antwerp, Belgium; Institut de Ciències del Mar (ICM), CSIC, 08003 Barcelona, Spain; Universidad Politécnica de Madrid (UPM) and Instituto Nacional de Investigación y Tecnología Agraria y Alimentaria (INIA-CSIC), Centro de Biotecnología y Genómica de Plantas, 28223 Madrid, Spain; University College Cork, APC Microbiome & School of Medicine, T12 K8AF Cork, Ireland

## Abstract

Autotrophic carbon fixation, the conversion of inorganic carbon into biomass, underpins life on Earth. Prokaryotes can carry out this process via at least seven biochemically distinct pathways, yet the phylogenetic and environmental distribution of most remains poorly resolved. Screening approximately 40 billion genes from reference genomes, metagenome-assembled genomes (MAGs) and unbinned metagenomic contigs, we provide a global assessment of the phylogeny and ecophysiology of prokaryotic autotrophs. Most pathway marker genes occurred in unbinned contigs and low-quality MAGs, representing phylogenetically distinct lineages absent from isolate genomes and quality filtered MAGs. Established autotrophs accounted for the large majority of pathway detections in quality filtered MAGs, largely recapitulating known biology from cultivated model organisms. Against this backdrop, the reductive TriCarboxylic Acid (rTCA) cycle, long considered restricted to anoxic environments, was detected in three phylogenetically distinct Campylobacterota lineages from oxygenated surface seawater, suggesting a previously unrecognized and unexpected niche for this pathway. We show that all three MAGs share an enzyme variant, previously described in other oxygen-tolerant lineages, that likely underlies their presence in the oxygenated surface ocean. Read mapping across global ocean metagenomes indicates that the organisms carrying it could be far more widespread than the scarcity of recovered MAGs alone would suggest. Together, these findings illustrate that the current view of global autotrophic carbon fixation is largely shaped by what genome-resolved methods can readily recover, while the true phylogenetic and ecological distribution of autotrophic carbon fixation appears to be much broader.

## Introduction

Autotrophic carbon fixation, the biochemical conversion of inorganic CO_2_ into organic biomass, is the foundation of primary production and a key driver of the global carbon cycle. Organisms capable of this process, autotrophs, synthesize their own organic carbon rather than acquiring it from other organisms, distinguishing them from heterotrophs that rely on organic carbon sources. Carbon fixation in a broader sense also encompasses non-autotrophic processes, such as anaplerotic CO_2_ assimilation in heterotrophic metabolism, which replenish metabolic intermediates rather than generate net new biomass from inorganic carbon (Erb, 2011). Throughout this study, we use “carbon fixation” to refer specifically to autotrophic processes.

The Calvin-Benson-Bassham (CBB) cycle, with Ribulose-1,5-bisphosphate carboxylase/oxidase (RuBisCO) as its central enzyme (Bassham & Calvin, 1960), is by far the most prominent carbon fixation pathway, present across cyanobacteria, many other autotrophic bacteria, and, through the endosymbiotic origin of the chloroplast, photosynthetic eukaryotes (Sagan, 1967; Tabita et al., 2008). Beyond the CBB cycle, at least six other biochemically distinct pathways have been described in prokaryotes (Hügler & Sievert, 2011; Ward & Shih, 2019). These include the reductive TriCarboxylic Acid (rTCA) cycle (Buchanan & Arnon, 1990; Evans et al., 1966), the Wood-Ljungdahl (WL) pathway (Ljungdahl, 1986), the reductive Glycine (rG) pathway (Figueroa et al., 2018; Sánchez-Andrea et al., 2020), the 3-HydroxyPropionate Bicycle (3-HPB, Herter, Fuchs, et al., 2002; Zarzycki et al., 2009), and the DiCarboxylate/4-HydroxyButyrate (DC/HB) and HydroxyPropionate/4-HydroxyButyrate (HP/HB) cycles used by autotrophic archaea (Berg et al., 2007; Huber et al., 2008; Könneke et al., 2014). Each pathway differs in energetic cost, oxygen tolerance, and phylogenetic and environmental distribution (Ward & Shih, 2019). While oxygenic photosynthesis using the CBB cycle dominates carbon fixation in sunlit environments, alternative pathways are known to contribute to carbon fixation across diverse environments, including the subsurface (Overholt et al., 2022; Rogers et al., 2023), soils (Su et al., 2026), marine sediments (Yue et al., 2026), oxygen minimum zones (Ruiz-Fernández et al., 2020), hydrothermal vents (Nakagawa & Takai, 2008), and stratified lakes (Alfreider et al., 2017). Even in sunlit surface waters, where oxygenic photosynthesis dominates, chemosynthetic carbon fixation has been estimated to contribute up to 22% of net primary production (Baltar & Herndl, 2019), yet which organisms and which pathways are responsible for this contribution remains largely unknown. Mapping the distribution of prokaryotic carbon fixation pathways across the Tree of Life and across environments is therefore a prerequisite for closing such gaps.

The rTCA cycle is one of the most ancient and phylogenetically widespread prokaryotic carbon fixation pathways (Becerra et al., 2014; Evans et al., 1966). It fixes an estimated 1 Pg of carbon annually, roughly 1% of the ∼100 Pg fixed via the CBB cycle, making it the second-largest contributor to global carbon fixation overall (Ward & Shih, 2019). Biochemically, the rTCA cycle shares most enzymes with the oxidative TCA cycle, the central catabolic pathway of aerobic respiration, but reductively fixes CO_2_ into acetyl-CoA rather than oxidizing acetyl-CoA to release energy. Three enzymatic steps cannot proceed in both directions and are replaced by dedicated enzymes in the rTCA cycle compared to its oxidative counterpart: ATP-citrate lyase cleaves citrate in place of citrate synthase, fumarate reductase reduces fumarate in place of succinate dehydrogenase, and 2-oxoglutarate:ferredoxin oxidoreductase (Oor) reductively carboxylates 2-oxoglutarate in place of the oxidative 2-oxoglutarate dehydrogenase complex (Buchanan & Arnon, 1990; Fuchs, 2011). A fourth enzyme, pyruvate:ferredoxin oxidoreductase (Por), is shared between both directions, oxidizing pyruvate to acetyl-CoA in many anaerobes and archaea or reductively carboxylating acetyl-CoA to pyruvate in the rTCA cycle, but is replaced by the irreversible pyruvate dehydrogenase complex in aerobes, which cannot support carbon fixation. Both oxidoreductases (Por and Oor) depend on iron-sulfur clusters that are sensitive to oxygen (Adams & Kletzin, 1996), and the stability of these clusters is thought to tightly constrain the rTCA cycle’s ecological distribution to anoxic and suboxic environments (Hügler & Sievert, 2011; Ward & Shih, 2019). Under this view, rTCA-based autotrophy is a strategy of specialized anaerobes and microaerophiles, fundamentally incompatible with fully oxygenated environments. Recent evidence, however, suggests this restriction may not be absolute. Oxygen-tolerant enzyme variants of Por and Oor have been described in a handful of aerobic and microaerophilic lineages spanning Aquificota and Campylobacterota (Bayer et al., 2021; Ikeda et al., 2006; Lücker et al., 2010; Molari et al., 2023; Yamamoto et al., 2003, 2006; Yun et al., 2002), and a broader genome survey showed that O_2_-sensitivity variation in these same enzyme families is present across Campylobacterota, Aquificota, and related phyla (Scott et al., 2026). Separately, read mapping against cultivated Arcobacter isolates has detected related sequences in surface ocean metagenomes (J. Li et al., 2024), though this detection has not been linked to the specific enzymatic adaptations that would allow oxygen-tolerant rTCA-based carbon fixation to actually occur there. These observations raise the possibility that oxygen-tolerant rTCA cycle variants extend beyond isolated lineages and contribute to rTCA-based carbon fixation beyond the pathway’s typical anoxic and microoxic niche, though linking such detection at global scale with the underlying genomic adaptation has so far been lacking.

Large integrated collections of reference genomes, global metagenomic datasets spanning diverse environments, and linked environmental data now make systematic characterization of prokaryotic diversity and functional capacities tractable at planetary scale (Fullam et al., 2023; Garritano et al., 2022; Kuhn et al., 2026; Nayfach et al., 2021; Richardson et al., 2023; Schmidt et al., 2024; Szabó et al., 2026). Yet genome-resolved approaches based on metagenome-assembled genomes (MAGs) capture only part of this diversity. Quality thresholds necessary for robust functional inference exclude a substantial fraction of environmental genomic content, with only 10% of assembled sequences in databases such as SPIRE meeting medium- or high-quality MAG criteria (Prasoodanan Pk et al., 2026; Schmidt et al., 2024), and the excluded, unbinned fraction harbors phylogenetically distinct lineages absent from assembled genomes altogether (Coelho et al., 2022; Prasoodanan Pk et al., 2026), suggesting that current surveys capture an incomplete picture of prokaryotic diversity and consequently their functional potential.

Here we present a large-scale survey of prokaryotic carbon fixation pathways across reference genomes, MAGs, and metagenomic contigs from global datasets (Fullam et al., 2023; Schmidt et al., 2024), based on Hidden Markov Models (HMMs) to detect marker genes for known carbon fixation pathways. This approach allowed us to identify genomes, MAGs, and contigs that carry the genomic potential for autotrophic carbon fixation, though the actual activity, and, in the case of the WL pathway and rG cycle, the direction in which the pathway operates, cannot be resolved from marker gene presence alone. Throughout this study, references to pathway presence or autotrophic genomes should therefore be understood as reflecting genomic potential inferred from marker gene detection, not confirmed metabolic activity. We first establish the phylogenetic and ecological distribution of all known carbon fixation pathways at this scale, then zoom into the rTCA cycle, hypothesizing that its restriction to anoxic and suboxic environments may be less absolute than currently assumed, and specifically test for its presence and persistence in the oxygenated and sunlit ocean. Our findings show that ecologically relevant functional diversity in unexpected environments is more widespread than previously appreciated, and that systematic inclusion of the unbinned genomic fraction will be essential for revealing its full extent.

## Results

### A curated marker gene pipeline reveals substantial phylogenetic diversity of carbon fixation marker genes beyond high-quality MAGs

We defined marker gene sets for seven carbon fixation pathways, with two and three sub-variants for HP/HB and rTCA respectively, yielding ten enzymatically distinct pathway variants in total. Because several pathways share enzymes with central metabolism or with each other, we distinguished these variants using required combinations of marker genes and, where applicable, co-occurrence rules, rather than relying on single diagnostic genes (Methods, Table S1). Using an HMM-based pipeline, we screened these marker sets across ∼40 billion genes from prokaryotic reference genomes (proGenomes3, Fullam et al., 2023) and MAGs and unbinned contigs (SPIRE, Schmidt et al., 2024) to assess the phylogenetic diversity and environmental distribution of marker genes for carbon fixation in global microbiomes.

We traced marker genes across four categories: reference genomes, medium/high-quality MAGs, low-quality MAGs, and unbinned contigs (Figure 1A). On average, 60.7% (range 23.2-92.8%) of all marker gene detections occurred exclusively in unbinned contigs or low-quality MAGs, fractions routinely excluded from downstream ecological analyses. To test whether this excluded fraction captures genuinely distinct lineages, rather than merely resampling diversity already present in the medium- to high-quality fraction, we performed rarefaction analysis (Figure 1B). Gene sequences were clustered at 95% identity to define species-level operational taxonomic units (Coelho et al., 2022). For the rTCA cycle, we excluded the citryl-CoA ligase (ccl) gene from rarefaction analyses, as it showed substantially higher detection rates than the co-occurring citryl-CoA synthase (ccsA and ccsB) genes, inconsistent with genuine pathway presence and more likely reflecting false positive hits. Rarefaction curves demonstrate that phylogenetic diversity increases substantially when unbinned contigs are included: even at the same number of samples, marker genes derived from all detection categories contain more species-level gene clusters than those from medium- and high-quality MAGs alone, with a median 1.51-fold increase across genes at matched sample size (range 1.02–2.80, Figure 1B). Terminal slopes of the rarefaction curves remained well above zero for both fractions (mean 0.198, range 0.04-0.49 for all detections versus mean 0.16, range 0.05-0.35 for medium- and high-quality MAGs alone, Figure 1C), indicating that diversity accumulation had not plateaued in either case. This suggests that the excluded fraction captures additional phylogenetic diversity not represented among assembled genomes, in line with recent analyses showing the unbinned fraction harbors a large share of uncharacterized phylogenetic diversity (Prasoodanan Pk et al., 2026).

**Figure 1.**
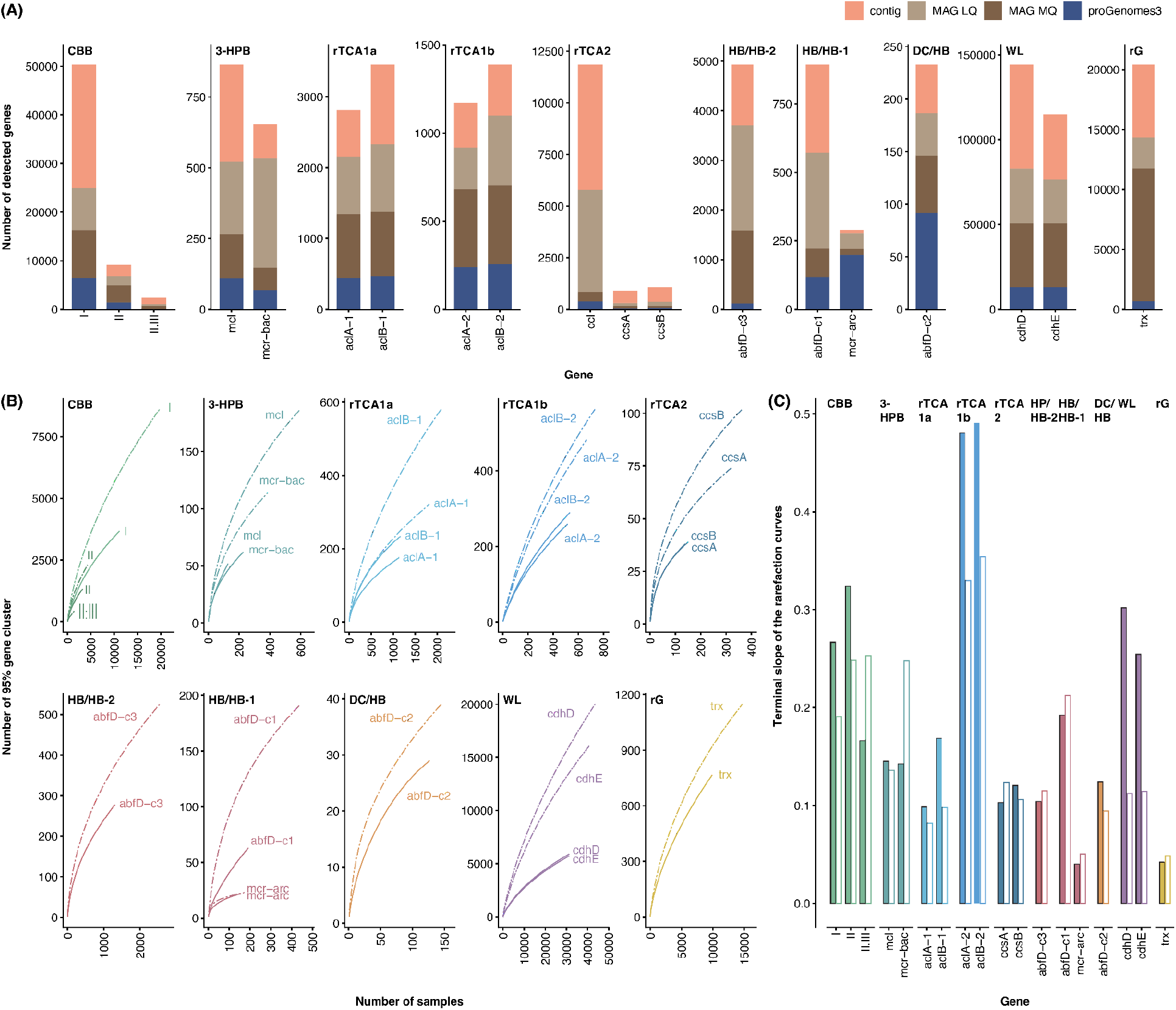
A genome-resolved marker gene pipeline reveals carbon fixation diversity beyond medium- and high-quality MAGs. (A) Detection of marker genes for ten carbon fixation pathway variants across reference genomes, medium/high-quality MAGs, low-quality MAGs, and unbinned contigs. (B) Rarefaction analysis comparing phylogenetic diversity of marker genes detected in reference genomes and medium-/high-quality MAGs (solid lines) versus all detections combined (dashed lines). (C) Terminal slopes of the rarefaction curves in (B), per gene, comparing the full detection (filled bars) set to reference genomes and medium-/high-quality MAGs alone (outlined bars).

### The distribution of carbon fixation pathways is tightly coupled to phylogeny and ecology across the prokaryotic tree of life

To resolve the phylogenetic, ecological and metabolic context of carbon fixation pathways, we integrated genomic, taxonomic and environmental metadata across 63,312 medium- and high-quality MAGs, providing sufficient genomic resolution for reliable taxonomic assignment and functional annotation. Habitat annotations were derived from Kim et al. and visualized by projecting samples onto their pre-existing taxonomic UMAP embedding (Kim et al., 2026, Table S2), while physiological traits were inferred using metaTraits (Podlesny et al., 2026, Methods). We then compared pathways to one another through enrichment analyses across habitat and physiology using two-sided Fisher’s exact tests with Benjamini-Hochberg correction for multiple testing (adjusted p < 0.05); only significant results are reported in the main text unless otherwise noted (Methods, Figure 2, Table S3).

**Figure 2.**
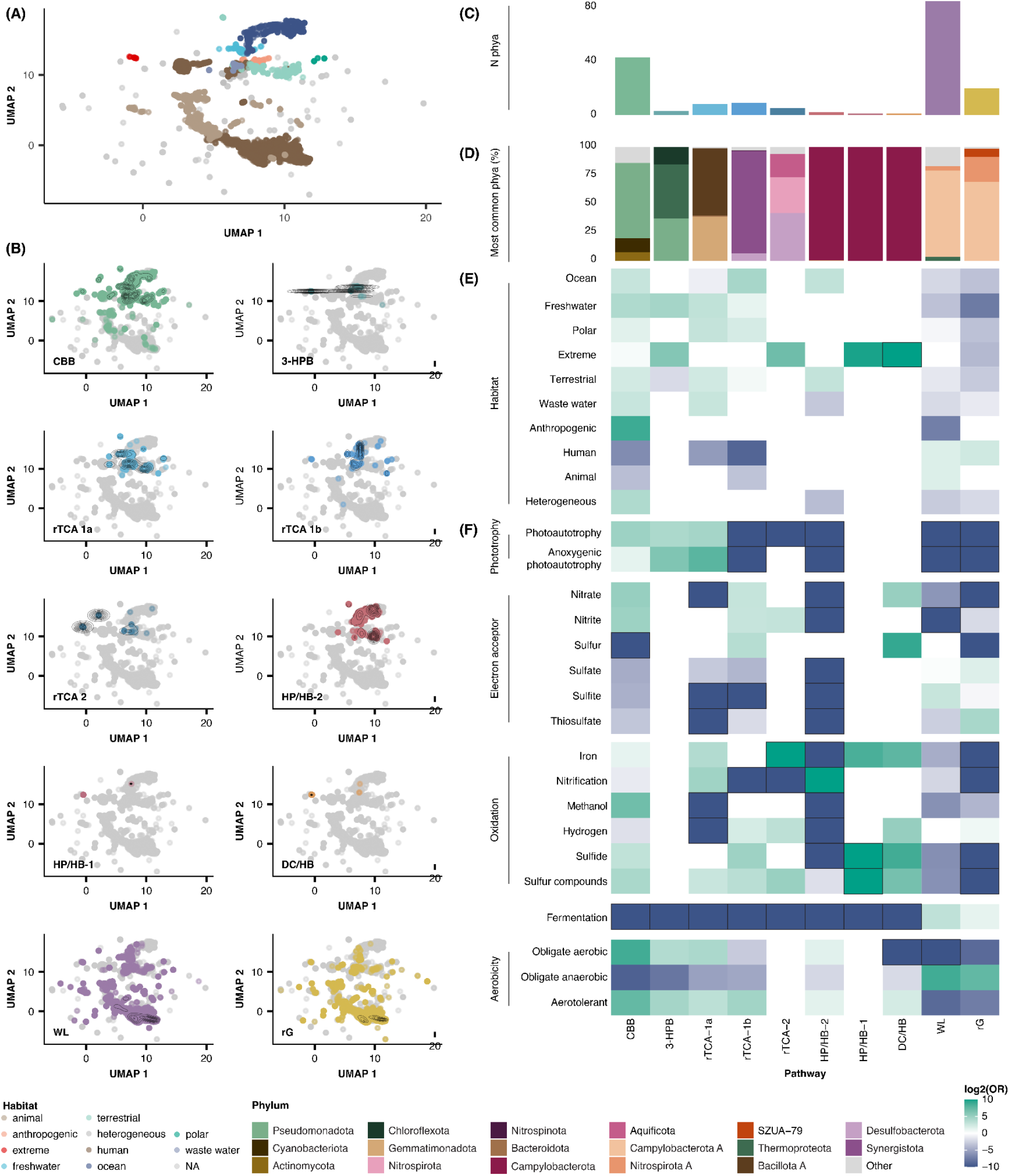
Carbon fixation pathway distribution is tightly coupled to phylogeny, habitat and metabolic strategy across the prokaryotic tree of life. (A) Microbiome samples positioned using the UMAP embedding from (Kim et al., 2026), with some categories combined, and colored by combined habitat type (Table S2). (B) The same UMAP embedding as in (A), with samples carrying each pathway highlighted. (C) Number of phyla in which each pathway was detected. (D) Percentage of MAGs belonging to the three dominant phyla for each pathway. (E) Odds ratios showing habitat enrichment for each pathway, based on the habitat classifications shown in (A). (F) Odds ratios showing functional enrichment for each pathway based on metaTraits annotations. For both (E) and (F), enrichment was computed by comparing all genomes carrying a given pathway against genomes carrying any other pathway (two-sided Fisher’s exact test, BH-corrected). Only significant enrichments are depicted in the heatmaps. This comparison is not adjusted for phylogenetic non-independence, which is addressed separately through phylum-resolved enrichment analyses (Methods, Table S3, S4).

In line with its known dominance in global carbon fixation, the CBB cycle was widespread phylogenetically and ecologically, detected across the 43 phyla and across all ten habitat categories, significantly enriched in eight of them relative to other pathways (OR up to 471.9). It was correspondingly enriched for diverse metabolic traits, including photoautotrophy (OR=28.0) alongside sulfur, iron, and methanol oxidation as well as nitrate reduction (OR range 2.3–120.8), matching the broad phylogenetic and metabolic diversity long associated with this pathway (Badger & Bek, 2008). Photoautotrophy was also enriched in the 3-HPB and rTCA1a cycle (OR=16.8 and 21.5), though biased towards the anoxygenic variant (OR=71.9 and 228.5). The rTCA1a cycle was additionally enriched for nitrification and iron oxidation (OR=25.2 and 14.1), and enriched in fresh water, polar, waste water, and terrestrial habitats (OR=5.0-8.5), while the 3-HPB, by contrast, showed no enrichment for other chemotrophic traits and was enriched only in extreme and freshwater habitats (OR=64.8 and 21.9). Unlike the rTCA1a cycle, the rTCA1b and rTCA2 cycle variants showed no phototrophic enrichment at all. The rTCA1b cycle nonetheless occupied a relatively broad niche, enriched for hydrogen, sulfur compounds oxidation, and nitrate reduction (OR=4.2-27.3). It was detected across six habitats but strongly enriched only in the ocean (OR=20.2). The niche of the rTCA2 cycle, in contrast, was more narrow. It was detected in only five phyla, confined almost entirely to extreme environments (OR=124.2), and strongly enriched for hydrogen, sulfur compounds, and iron oxidation (OR=9.6-5173.4), more consistent with the classical framing of the rTCA cycle as thermophilic chemolithotrophic pathway (Hügler & Sievert, 2011). DC/HB and HP/HB-1 showed a similarly narrow distribution, restricted almost exclusively to Thermoproteota in extreme environments and strongly enriched for iron and sulfur-compound oxidation (DC/HB: OR=100.1-561.5; HP/HB-1: OR=348.5-4226.0), paralleling their known role as chemolithotrophic carbon fixers in hydrothermal and geothermal systems (Berg et al., 2007; Huber et al., 2008). The HP/HB-2 cycle diverged sharply from this pattern despite also being detected almost exclusively in Thermoproteota, and within them, the class Nitrososphaeria specifically. They were enriched instead for nitrification and obligate aerobicity (OR=16384.0 and 2.8) and for terrestrial and ocean habitats (OR=7.9 and 8.4), matching the known ecology of Nitrososphaeria as globally distributed ammonia-oxidizing archaea (Könneke et al., 2014; Stahl & de la Torre, 2012). The WL pathway was the phylogenetically most widespread pathway and was detected in 85 phyla and the rG cycle was detected in 20 phyla and both were detected across most habitats, reflecting their broad taxonomic distribution among anaerobes (Ragsdale & Pierce, 2008; Sánchez-Andrea et al., 2020). Unlike the CBB cycle, however, this broad phylogenetic distribution did not result in a correspondingly diverse ecological niche for either. Both were significantly enriched only in human- and animal-associated habitats (OR=3.3-5.2), and correspondingly enriched strongly for obligate anaerobicity (OR=428.0 and 179.4).

These pathway-level signatures often masked divergent strategies among the major phyla carrying each pathway. To resolve this, we performed Fisher’s tests again within each pathway, comparing MAGs from each of the most common phyla against all other MAGs carrying the same pathway, revealing that pathways with a shared overall enrichment frequently comprised phyla with distinct, sometimes opposing, individual strategies (Methods, Table S4). Among MAGs carrying the CBB cycle, only those belonging to Cyanobacteriota were significantly enriched for photoautotrophy, and specifically enriched in polar lakes and freshwater habitats (OR=10.5 and 4.4), in concordance with their established role as oxygenic photosynthesizers (Flombaum et al., 2013). The numerically dominant Pseudomonadota was instead significantly enriched for chemolithotrophy through sulfur oxidation and for anthropogenic habitats (OR=21.0 and 7.6), and the third dominant phylum, Actinomycetota, was significantly enriched for fermentation and obligate anaerobicity relative to the other two (OR=8.0 and 1.9), matching this phylum’s broader association with soil and host-associated lifestyles (Ventura et al., 2007). The three rTCA cycle variants each showed similar differences between phyla. Within rTCA1a, nitrifying Nitrospirota were enriched in terrestrial and waste water habitats (OR=13.5 and 47.0), matching their established role in nitrification across engineered and soil systems (Freitag et al., 2005; Gruber-Dorninger et al., 2015; Spieck et al., 2006), while anoxygenic phototrophic Bacteroidota were instead enriched in freshwater (OR=52.1), matching the known ecology of green sulfur bacteria in stratified freshwater systems (S. L. Garcia et al., 2021). Within rTCA1b, the dominant Campylobacterota carried the pathway’s broad marine distribution, though within-pathway comparison identified them specifically enriched in terrestrial habitats (OR=5.5), while the Aquificota lineage was instead enriched for extreme habitats and hydrogen oxidation (OR=55.1 and 27.3). Although the rTCA2 cycle was confined to a single narrow niche overall, with all three of its major phyla sharing the capacity for iron oxidation, they still showed diverging metabolic properties: Nitrospirota_A were uniquely enriched for obligate aerobicity, Aquificota uniquely for obligate anaerobicity and additionally for sulfur-compound oxidation (OR=17.9), and SZUA-79 uniquely for sulfate reduction. Also the WL pathway and rG cycle show a similar pattern. The dominant Bacillota_A were significantly enriched for fermentation in both (OR=4718.3 and 9620.6), while a smaller group of Desulfobacterota was instead independently enriched for sulfate and thiosulfate reduction (OR up to 407.9). A third WL-associated phylum, Chloroflexota, was additionally enriched for iron oxidation (OR=18.2) and, atypically for this pathway, obligate aerobicity (OR=84.3), a result we consider likely to reflect an artifact given it is based on a small percentage of genomes carrying this trait (1.7%) though it nonetheless represents a significant enrichment compared the overall prevalence across the pathways is much lower (0.05%).

Together, these results provide a global survey of prokaryotic autotrophy, resolving their phylogenetic, environmental and metabolic niches. Each pathway occupied its own distinct niche, and where several phyla carried the same pathway, they frequently contributed diverging, sometimes opposing, ecophysiological profiles rather than a single shared strategy.

### rTCA-carrying Campylobacterota recur in the oxygenated surface ocean

Among the pathways surveyed, the rTCA cycle stood out for a departure from the common framing of this pathway as restricted to anoxic and microoxic conditions (Hügler & Sievert, 2011). While the WL pathway and rG cycle were both enriched for obligate anaerobicity, the three rTCA variants showed no such signature. Obligate aerobicity or aerotolerance were instead detected across several rTCA-carrying phyla (Table S4, Note S1), including Nitrospirota (95.6% obligate aerobic) and Campylobacterota (89.6% aerotolerant). Nitrospirota’s aerobicity matches their known role as nitrifiers, whereas Campylobacterota’s aerotolerant pattern may reflect adaptation to fluctuating, rather than persistently anoxic, redox conditions, characteristic of the environments these lineages typically inhabit (Glaubitz et al., 2009; Grote et al., 2008). A recent description of a *Sulfurimonas* lineage, *S. pluma*, thriving in fully oxygen-saturated hydrothermal plume water challenges this interpretation, suggesting that at least some Campylobacterota are adapted to fully oxic conditions rather than merely tolerant of fluctuation (Molari et al., 2023), and raising the question of whether this capacity extends to other oxygenated marine habitats, including the surface ocean. We therefore examined the distribution of the 196 marine Campylobacterota MAGs in our dataset across marine habitats and oxygen concentrations.

Most marine Campylobacterota MAGs carrying the rTCA cycle for which oxygen data were available came from low-oxygen or microaerobic settings, reflecting the pathway’s classical association with low oxygen (Table S5). Two, however, were recovered from oxygenated waters: MAG IO (Sulfurimonadaceae), off Madagascar at 7 m and 217 µmol/L O_2_, and MAG HL (Sulfurovaceae), off Helgoland in the North Sea at 1 m and 296 µmol/L O_2_. Both oxygen values were inferred from World Ocean Atlas climatology (H. E. Garcia et al., 2024), but given the shallow depth of both samples, where climatological estimates closely track actual conditions, these values likely reflect the true oxygen environment. In addition, community profiles (mOTU relative abundance, Supp. Data) were dominated by classic oxic surface-ocean taxa, *Prochlorococcus* and *Synechococcus* at the Madagascar site, and SAR11/Pelagibacteraceae at both sites, independently supporting the oxic classification. A third Campylobacterota MAG carrying marker genes for the rTCA cycle was independently detected in the MAG catalogue from a 15-year time series at 20 m depth from the Blanes Bay Microbial Observatory (Latorre et al., 2025, Figure 3, MAG BB, Arcobacteraceae) at a median relative abundance of approximately 0.001% across all 175 samples spanning the full 15-year period (detected in 175/175 samples; range 6.4×10⁻⁸–2.6×10⁻³%), placing it among the least abundant 16% of the 1,505 genomes profiled at this site (Latorre et al., 2025).

**Figure 3.**
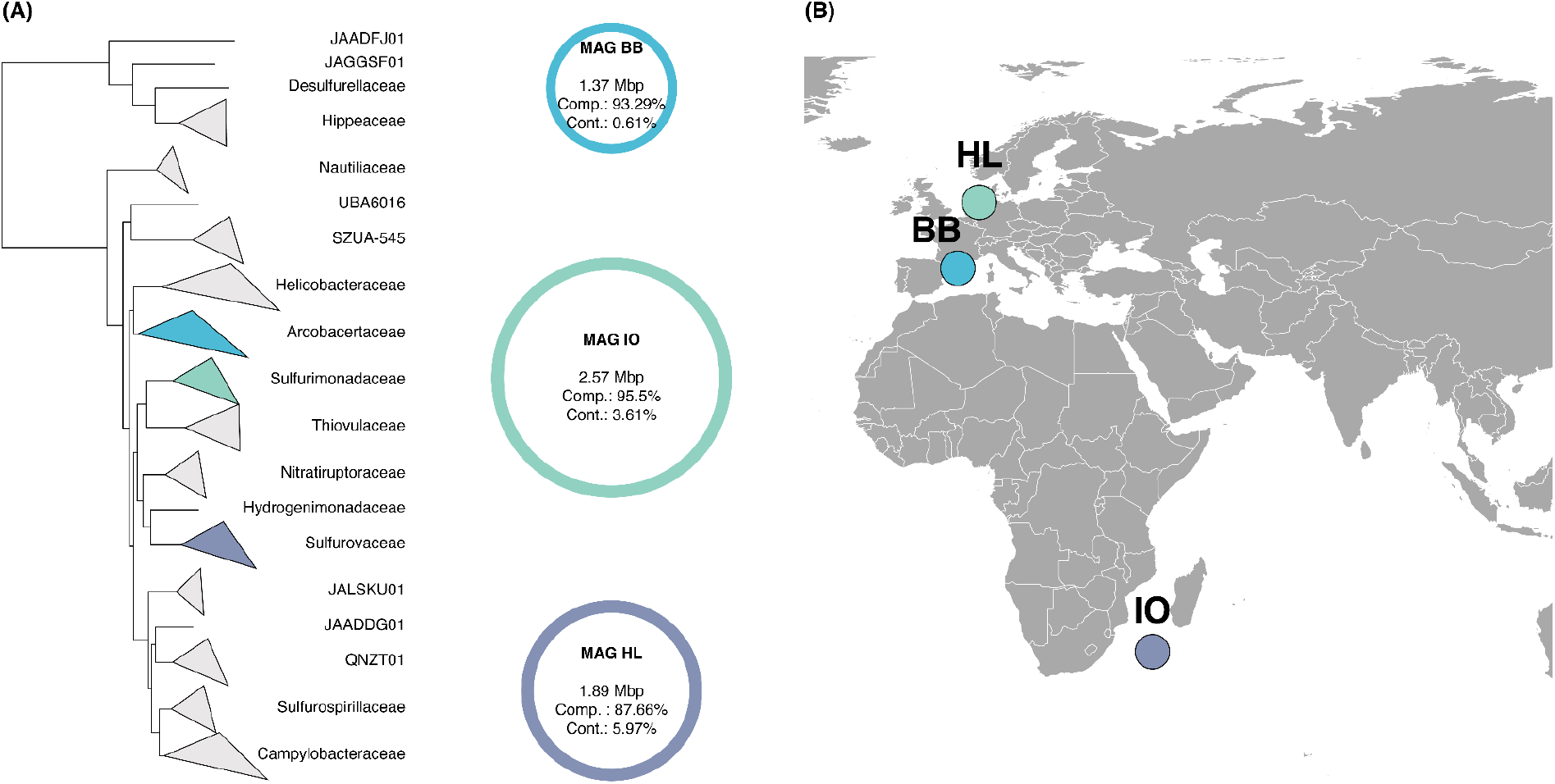
rTCA-carrying Campylobacterota occupy oxygenated surface waters across phylogenetically distinct lineages. (A) Phylogeny of Campylobacterota (based on the GTDB r220 tree), with families collapsed and the three families containing MAG BB, MAG HL, and MAG IO highlighted in MAG-specific colours. Next to the tree, each MAG is represented by an unfilled circle outlined in its corresponding colour and scaled in size to genome size (Mbp), with MAG name, genome size (Mbp), completeness (Comp.), and contamination (Cont.) labelled inside. (B) Sampling locations of the three Campylobacterota MAGs in the surface ocean.

### Comparative genomics identifies a shared enzyme variant associated with oxygen tolerance in rTCA-carrying Campylobacterota

To characterize the three rTCA carrying MAGs from the surface ocean, we performed functional annotations of their genomes. These indicated that all three encode the marker genes for the rTCA cycle and appear capable of using hydrogen and sulfur oxidation as electron donors, mirroring the broader metabolic profile of autotrophic Campylobacterota (Note S2, Table S6, S7). Their presence in the surface ocean raised the question whether these MAGs share any of the adaptations that allow other related lineages to persist in oxygenated habitats. We thus compared the three MAGs, MAG HL, MAG BB, and MAG IO, with two MAGs of *S. pluma*, the lineage with characterized adaptations to oxygenated hydrothermal plumes, including oxygen-tolerant enzyme variants and high-oxygen-affinity respiration (Molari et al., 2023) and with reference genomes from proGenomes3 (Fullam et al., 2023) sharing the same GTDB r220 genus annotations as our MAGs of interest, to establish whether the features observed in our three MAGs are typical within their respective genera (Figure 2, Table S8, Parks et al., 2022).

One adaptation detected in *S. pluma* is a shift in respiratory strategy, using oxygen, rather than nitrate, as the terminal electron acceptor. This is reflected in the loss of nitrate respiration genes and the acquisition of a caa₃-type oxidase. Unlike the cbb₃-type oxidase typical of Campylobacterota (Han & Perner, 2015; Labrenz et al., 2013; Molari et al., 2023), caa₃ supports efficient respiration under permanently high oxygen (Pitcher & Watmough, 2004; Sousa et al., 2012). MAG HL and MAG IO match this strategy. Both encode an additional caa₃-type oxidase and largely lack nitrate respiration genes (napAB absent in HL, napA absent in IO), contrasting with their respective genera, where cbb₃-type oxidase and denitrification predominate (Figure 2, Table S7, Table S8). MAG BB does not share this adaptation. Like all other Arcobacteraceae, it retains genes for nitrate reduction (napAB present) alongside cbb₃-type respiration, supplemented by a bd-type oxidase (Table S7). MAG BB’s presence in oxygenated surface water despite lacking the caa₃-based specialization indicates that this respiratory adaptation is not strictly required for rTCA-cycle-carrying Campylobacterota to occur in oxygenated water.

In contrast to their divergent respiratory profiles, all three MAGs share a common enzyme variant in the rTCA cycle’s oxygen-sensitive carboxylating enzymes, pyruvate:ferredoxin oxidoreductase (Por) and 2-oxoglutarate:ferredoxin oxidoreductase (Oor), homologous to characterized oxygen-tolerant forms. These oxygen-tolerant variants of both enzymes, containing five subunits rather than the four typical of Campylobacterota, were first described in *Hydrogenobacter thermophilus* (Ikeda et al., 2006; Yamamoto et al., 2003, 2006; Yun et al., 2002) and later shown by Molari et al. to also occur in the oxygen-tolerant *S. pluma* (Molari et al., 2023). Through phylogenetic placement of the alpha subunits and confirmation of the full five-gene operon (Methods), we found that all three surface ocean MAGs likewise encode this five-subunit variant of both enzymes. Phylogenetically, these variants place together with those of *S. pluma* and separately from the four-subunit form, which the majority of other Campylobacterota references retained. The five-subunit form was nonetheless present in a subset of reference genomes as well, occurring sporadically across genera rather than being confined to a single representative species (Figure 4B, Figure S1, Table S8).

**Figure 4.**
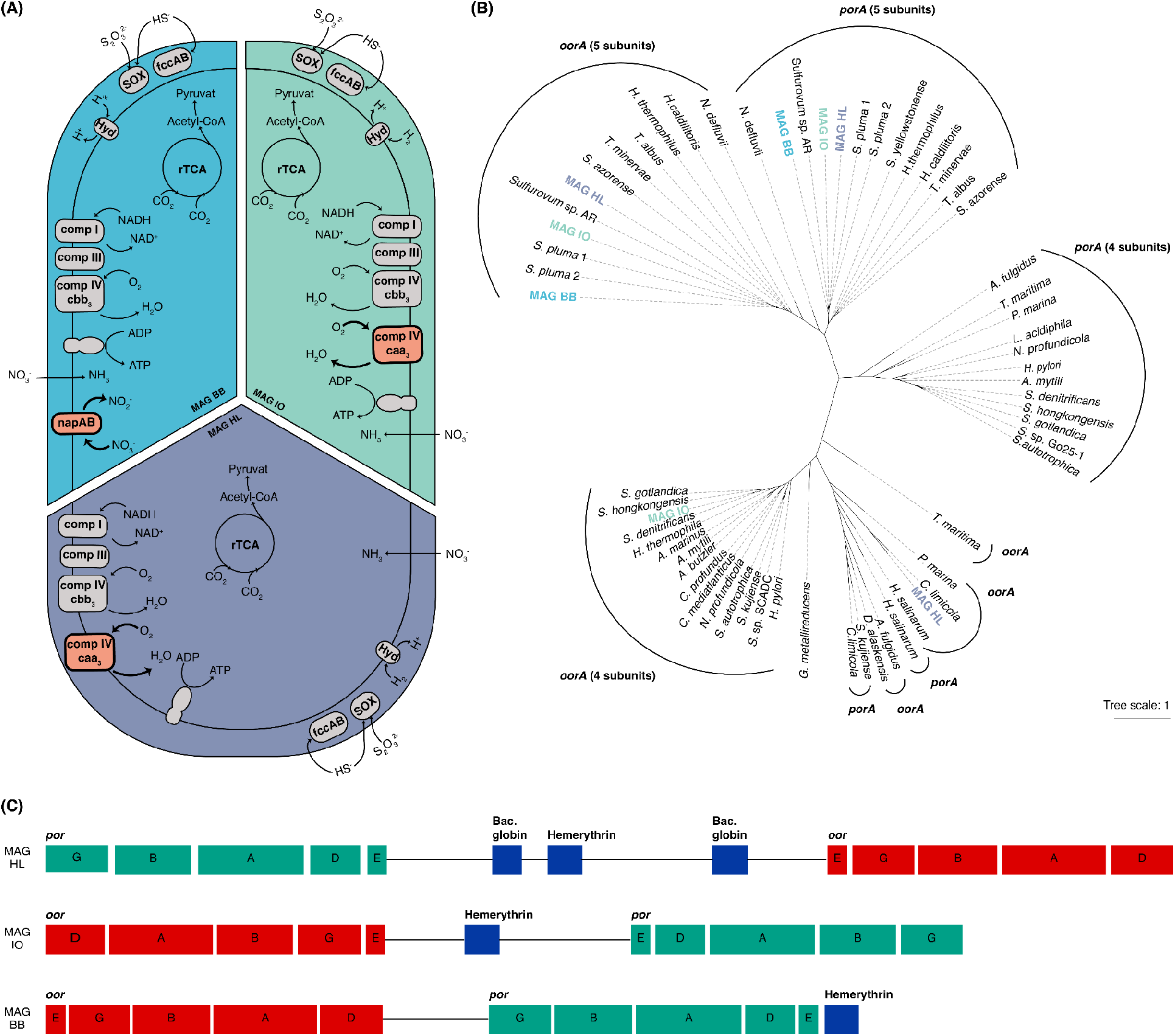
A shared five-subunit enzyme variant, rather than respiratory strategy, appears to underpin the presence of three rTCA-cycle-carrying Campylobacterota lineages in the oxygenated surface ocean. (A) Schematic summary of metabolic profiles for the three MAGs, shown as an oval cell diagram divided into MAG-specific sections, with differences in the respiratory profile highlighted in orange (Note S2, Table S6, S7). (B) Maximum-likelihood phylogeny of the alpha subunit of the Por/Oor, including sequences from the three MAGs and published reference sequences (UniProt), including *H. thermophilus* and S. *pluma*. Four- and five-subunit variants were distinguished by phylogenetic placement of the alpha subunit and confirmation of the complete five-gene operon in the corresponding genomic neighbourhood (Figure 4C, Methods). The extended tree including genus-matched proGenomes3 reference genomes is shown in Figure S2 (Table S8). (C) Genomic organization of the por and oor operons in the three MAGs, showing the arrangement of individual subunits and the co-localization of bacterial globin and hemerythrin genes on the same contigs (Table S6).

The genomic organization of these enzymes further points to co-regulation or a shared evolutionary origin of these gene clusters. In all three MAGs, the Por and Oor operons occur in close proximity, matching the organization seen in *S. pluma* and Aquificota (Figure 4C) whereas in reference genomes carrying the canonical four-subunit form these genes dispersed more broadly (Table S8). In addition, five-subunit Por and Oor genes are clustered with genes encoding hemerythrin and bacterial globins, which may play a role in oxygen detection, storage, and detoxification (Figure 4C, French et al., 2008; Kendall et al., 2014). This co-localization suggests an integrated module pairing enzyme isoforms associated with oxygen tolerance with local oxygen-scavenging genes.

### Populations related to the oxygen-tolerant Campylobacterota lineages recur across the surface ocean

The three MAGs recovered here may represent only a small fraction of where these oxygen-tolerant lineages actually occur. To test this, we mapped metagenomic reads from the TARA Oceans dataset against the three MAGs (Karsenti et al., 2011), counting a MAG as detected when at least 50% of its genome length was covered by mapped reads at 95% identity. Detection frequencies differed sharply: MAG BB was detected in 501 of 964 screened samples, indicating populations closely related to this lineage are widespread rather than locally restricted. MAG HL was detected in fewer samples overall (43) yet across ocean regions, and MAG IO in only two from the Pacific. All three were detected predominantly in the surface ocean, with slightly diverging distributions across depth, oxygen, and nitrate. MAG BB spanned the full oxygen gradient (0.6–393 µmol/L O_2_) and nitrate up to 44 µmol/L, while MAG HL and MAG IO were confined to higher-oxygen, lower-nitrate samples (Figure 5), matching the differing respiratory strategies observed in the reference MAGs themselves, though whole-genome mapping alone cannot confirm that the environmental populations detected here share these same respiratory adaptations. Neither genome coverage nor sequencing depth showed a clear association with oxygen, nitrate, chlorophyll, temperature, or salinity for MAG BB and MAG HL (Table S9), indicating they occur across a broad range of productivity and nutrient regimes. Matching metatranscriptomic samples showed the MAGs transcriptionally active in the large majority of detected samples (≥10 genes transcribed: 78% MAG BB, n=501; 77% MAG HL, n=43; 100% MAG IO, n=2), with no significant differences in depth, oxygen, nitrate, or geography between samples showing transcriptional activity and the full set of detected samples (Figure 5). We additionally screened for the rTCA marker genes (aclA/aclB, por and oor subunits) across the same samples. Detection was markedly sparser than for the whole genome (recovered in only 10.4% and 9.3% of genome-detected samples for MAG BB and MAG HL, respectively, and in none for MAG IO), possibly reflecting low and uneven coverage caused by these organisms’ low abundance (Supp. Data). Regardless, these lineages are themselves widely distributed and active across the surface ocean, a pattern worth exploring further to understand where and how broadly the oxygen-tolerant rTCA cycle variants occur.

**Figure 5.**
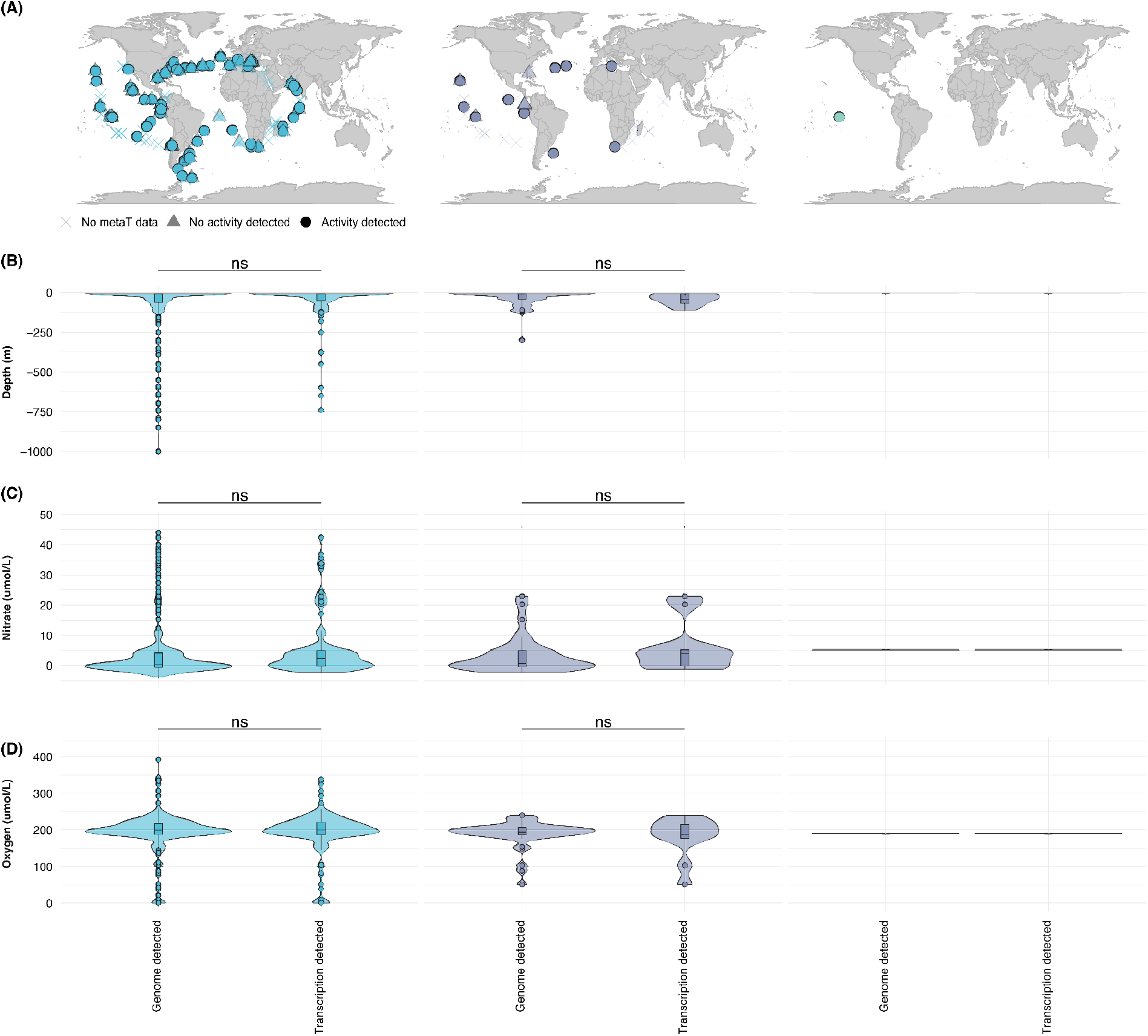
Campylobacterota lineages related to the rTCA-carrying MAGs are globally distributed across the surface ocean. (A) Geographic distribution of TARA Oceans samples in which MAG BB, MAG HL, and MAG IO were detected via metagenomic read mapping (≥50% genome coverage), faceted and coloured by MAG. Point shape indicates whether the sample also met the metatranscriptomic activity threshold (circle: ≥10 genes transcribed; triangle: below threshold or no metatranscriptomic data available). (B–D) Box- and violin plots showing depth (B), oxygen concentration (C), and nitrate concentration (D) at TARA Oceans sampling sites where each MAG was detected. Each panel is faceted and coloured by MAG, with samples further split by whether they were only detected in the metagenomic or also the metatranscriptomic data.

## Discussion

The rTCA cycle has long been considered restricted to anoxic and suboxic environments, its ecological range defined by the oxygen sensitivity of its core carboxylating enzymes (Hügler & Sievert, 2011; Ward & Shih, 2019). Our global metagenomic survey identified three phylogenetically distinct Campylobacterota lineages carrying the rTCA cycle in oxygenated surface waters. Read-based screening of the TARA Oceans dataset showed that these lineages are not confined to the handful of sites where they were originally recovered: across approximately 964 samples spanning multiple ocean regions, MAG BB alone was detected in around 501 samples and MAG HL in 43, the large majority of which were also transcriptionally active. Combined with a 15-year coastal time series showing one of these lineages persisting at low but stable abundance, these results indicate that rTCA-carrying Campylobacterota are rare but recurrent across the contemporary, oxygenated surface ocean, rather than an anomalous occurrence. This raises a direct mechanistic question: how do organisms employing an inherently oxygen-sensitive pathway persist in oxygenated waters? Prior work has described two distinct solutions to this problem. One solution, detected in epibionts of deep-sea mussels, is the acquisition of a complete CBB cycle via multiple HGT events from co-occurring symbionts, replacing their ancestral rTCA machinery entirely as an evolutionary response to oxygen exposure in fluctuating redox conditions (Assié et al., 2020). The other is to modify the rTCA cycle’s core carboxylating enzymes directly. Oxygen-tolerant variants consisting of five subunits of both pyruvate:ferredoxin oxidoreductase (Por) and 2-oxoglutarate:ferredoxin oxidoreductase (Oor), were first described in *H. thermophilus* (Ikeda et al., 2006; Yamamoto et al., 2003, 2006; Yun et al., 2002) and have since been found in aerobic nitrifiers (Bayer et al., 2021; Lücker et al., 2010) and in the oxygen-adapted hydrothermal-plume species *S. pluma* (Molari et al., 2023). A recent genome survey across autotrophic Bacteria further shows that O2-sensitivity variation in these same enzyme families broadly tracks oxygen lifestyle across Campylobacterota, Aquificota, and related phyla (Scott et al., 2026), suggesting this kind of enzymatic adaptation may be more widespread across the rTCA cycle than previously appreciated, though direct evidence for this breadth has so far come only from reference genomes and cultured isolates. Our findings extend evidence for these enzyme variants into environmental metagenomic samples and show that rTCA-cycle-carrying organisms can occur in fully oxygenated habitats. All three MAGs described here carry the same five-subunit Por/Oor variant, and were detected across surface waters from three different ocean regions and three distinct Campylobacterota families, showing that this enzyme variant is not confined to *S. pluma*, and indicating it is more broadly distributed across marine Campylobacterota than previously known. Phylogenetic reconstruction placed all Campylobacterota five-subunit variants within a single monophyletic clade sister to Aquificota, suggesting acquisition via horizontal transfer from an Aquificota-related donor. Within Campylobacterota, however, genomes carrying the five subunit variants did not cluster by genus or family. Instead, five subunit variants were present only sporadically among close relatives within the same genus. This distribution is more consistent with subsequent horizontal spread among Campylobacterota lineages than with simple vertical inheritance following a single ancestral acquisition.

The recovery of surface ocean MAGs carrying the rTCA cycle from only three sites despite global sampling, together with the stable median relative abundance of approximately 0.001% maintained by MAG BB across 15 years at Blanes Bay, and the uniformly low genome coverage across all three MAGs in TARA Oceans metagenomes, place the Campylobacterota identified here in the rare biosphere. Rare-biosphere taxa have in some cases been shown to exert disproportionate biogeochemical influence elsewhere (Jousset et al., 2017; Sogin et al., 2006). This rarity likely explains why such organisms have been previously overlooked, despite their potential ecological relevance. Campylobacterota are already established as quantitatively important autotrophs elsewhere, dominating carbon fixation at pelagic chemoclines in the Baltic and Black Seas (Glaubitz et al., 2009; Grote et al., 2008) and contributing measurable autotrophic activity across Arcobacter, Sulfurimonas and Sulfurovum lineages at hydrothermal vents (McNichol et al., 2022). Whether they contribute similarly in the oxygenated surface ocean, where conditions differ fundamentally from chemoclines and vents, cannot be assumed from these systems alone and remains to be tested directly. Chemosynthetic carbon fixation has separately been estimated to contribute up to 22% of net primary production in surface waters (Baltar & Herndl, 2019), yet the organisms responsible remain poorly resolved, and the Campylobacterota identified here are candidates to fill part of that gap. Biochemical characterization of the five-subunit enzyme variants identified here, targeted activity measurements across defined redox gradients, and expanded geographic and temporal sampling will each test and extend the specific predictions this survey generates, and closer attention to the unbinned fraction may show how far this kind of overlooked diversity extends across microbial carbon fixation more broadly.

The discovery of the three rTCA carrying MAGs in the surface ocean contrasts a broader pattern established in this survey. Across the ten pathways variants characterized, the phyla, habitats and metabolic capacities we could assign to the pathways based on MAG annotations largely recapitulated previous findings, including Cyanobacteriota as oxygenic photoautotrophs (Flombaum et al., 2013), Nitrospirota as nitrifiers (Bayer et al., 2021; Lücker et al., 2010), and Thermoproteota as either thermophilic chemolithotrophs or ammonia oxidizers (Berg et al., 2007; Huber et al., 2008; Könneke et al., 2014; Stahl & de la Torre, 2012). This reflects how genome-resolved surveys work. Current standards for metagenomic analysis recommend limiting downstream functional inference to medium- and high-quality MAGs (Bowers et al., 2017), an approach that favors organisms abundant enough to assemble well and, as a result, organisms that are often already well characterized (Parks et al., 2017). Despite this limitation, we observed another interesting case that warrants further investigation. While 3-HPB is well studied in Chloroflexota, known anoxygenic photoautotrophs from microbial mats in hot springs, its putative role in other organisms and habitats is not well established (Klatt et al., 2007). We also detected 3-HPB in Pseudomonadota and Gemmatimonadota, mostly from freshwater samples, which showed no enrichment for photoautotrophy or for any chemoautotrophic pathways tested. Freshwater Pseudomonadota and Gemmatimonadota have been reported as obligate photoheterotrophic members of the aerobic anoxygenic phototrophic community (Piwosz et al., 2022; Villena-Alemany et al., 2023, 2024), but the presence of 3-HB marker genes in these organisms, both in our data and in previous reports (Garritano et al., 2022; Kleiner et al., 2012), suggests they may represent yet undiscovered players in alternative carbon fixation, though in this specific case full characterization and validation will be needed to confirm their role.

Our observation that MAG-based characterization only sporadically reveals pathways associated with unexpected organisms or niches is underpinned by a larger structural limitation of MAG-based surveys. In our survey, 60.7% of all marker gene detections occurred exclusively in unbinned contigs or low-quality MAGs. Because assembly and binning both rely on sequencing coverage, this fraction is disproportionately populated by rare, low-abundance organisms that could not be assembled into a usable genome at all. This creates a systematic split where MAG-based studies preferentially recover abundant organisms, which are also more likely to have already been described, while novel findings are more often confined to the low-quality and unbinned fraction, where limited genomic completeness and taxonomic resolution leave relatively little room to characterize them further. Rarefaction analyses show that this split does not merely result in a difference of absolute detections, but that the low-quality and unbinned fraction contains genetic and potentially functional variants not found among medium- and high-quality MAGs at all, matching recent work showing that this fraction harbors phylogenetically distinct lineages absent from assembled genomes entirely (Prasoodanan Pk et al., 2026). Fully exploiting the much larger unbinned fraction for functional inference remains a methodological challenge and cannot yet provide comprehensive mechanistic or ecological insight, given limited genomic completeness and taxonomic resolution. We therefore treat it not as a source of robust functional claims in its own right, but as a signal pointing toward where future discovery is most likely.

This survey provides a tree-of-life-wide characterization of how prokaryotic carbon fixation pathways are distributed, showing that its phylogenetic, ecological, and metabolic distribution is tightly coupled across all three. Despite the systematic bias toward abundant and well characterized organisms this approach entails, it also revealed a known pathway in a previously uncharacterized setting: three phylogenetically distinct Campylobacterota lineages carrying the rTCA cycle in oxygenated surface waters, a habitat long considered fundamentally incompatible with this pathway. A shared enzyme variant, rather than any particular respiratory strategy, is most likely what enables these organisms to occur there. Genomic and transcriptomic detections suggest they recur across the surface ocean rather than being confined to the three MAGs described here. Together, these findings point to a potentially ecologically relevant mode of carbon fixation that had gone largely unrecognized, and to how much more likely remains to be found in the majority of environmental sequence data that current methods still cannot resolve.

## Methods

### Selection of marker genes

Marker gene selection for each carbon fixation pathway was primarily based on the METABOLIC annotation framework (Zhou et al., 2022) and the selection described by (Jaffe et al., 2023), with minor adjustments described below (Table S1).

For the CBB cycle, Form I and Form II RuBisCO large subunits were used, as this enzyme is the pathway’s defining enzyme, with no equivalent role in any other carbon fixation pathway. Form III and Form IV RuBisCO were included as false positive controls, as their roles are unclear and their presence is not considered a reliable indicator of CBB cycle activity. For the 3-HPB, malonyl-CoA reductase/3-hydroxypropionate dehydrogenase (mcr-bac, K14468) and malyl-CoA lyase (mcl, K08691) were used. We used mcl instead of propionyl-CoA synthase used in other studies (Jaffe et al., 2023; Zhou et al., 2022), as it is a well-characterized, bifunctional enzyme central to this pathway (Herter, Busch, et al., 2002), and together with mcr-bac, which is not shared with other pathways, provides a reliable basis for pathway presence. For the HP/HB cycle variants and DC/HB cycle, we used the 5-hydroxybutyryl-CoA dehydratase (abfD) gene, as it is a shared marker across all three pathway variants, distinguishing between them through phylogenetic identity (Figure S10) and complemented detection of the classical, Crenachaeae variant of the HP/HB cycle with the homologue malonyl-CoA reductase gene (KO K15017), which is not part of the DC/HB cycle and lacking/replaced by a distinct homologue in the HP/HB-2 variant (Könneke et al., 2014; F. Li et al., 2018). For the WL pathway, solely the CO-dehydrogenase/acetyl-CoA synthase subunits cdhD (K00194) and cdhE (K00197) were used, while cdhA/cooS (K00198) was excluded, as it shares homology to other carbon monoxide dehydrogenases which are not specific to the pathway (Adam et al., 2018). For the rTCA cycle, use distinct enzymes performing the citrate cleavage step and distinguish different pathway variants (Hügler & Sievert, 2011): we used two variants each of the ATP citrate lyase subunits aclA and aclB (K15230, K15231) that distinguish the rTCA1a and rTCA1b cycle variants, and citryl-CoA synthetase/lyase (ccsA, ccsB, ccl) which is specific to the rTCA2 cycle variant. For the rG pathway, there was no established blueprint to build on, as it is often omitted from such surveys altogether (Garritano et al., 2022; Jaffe et al., 2023; Zhou et al., 2022), and no comprehensively validated marker gene set currently exists for it. We used thioredoxin reductase (trx) which is part of the glycine reductase complex (Sánchez-Andrea et al., 2020) in place of the more pathway-specific glycine cleavage/synthase system, which had too few sequences represented in the reference database to support reliable competitive detection at the scale of this survey. rG-associated trx sequences from organisms characterized by (Sánchez-Andrea et al., 2020) form a phylogenetically distinct clade, and marker gene searches were restricted to this clade specifically rather than to trx broadly, which is not itself pathway-specific. This restriction was tested against a reference set of 10,182 *Escherichia* genomes present in SPIRE (Schmidt et al., 2024), in which trx is present but the reductive glycine pathway is not, and returned a false-positive rate of 0.48% (49/10,182).

### RuBisCO reference phylogeny and profile HMM construction

RuBisCO reference sequences were obtained from proGenomes3 (Fullam et al., 2023). Predicted proteins were screened against a Pfam domain PF00016 using hmmsearch (HMMER, v3.3.1, Eddy, 2011). Resulting hits were clustered at 99% identity using MMseqs2 (commit ca58693; mmseqs cluster, default parameters, --min-seq-id 0.99, (Steinegger & Söding, 2017), and sequences shorter than 100 or longer than 750 amino acids were removed. Reference sequences from (Erb et al., 2012) were added to the dataset (333 sequences in total). Sequences were aligned using FAMSA v1.6.1 (Deorowicz et al., 2016) and trimmed using trimAl (v1.4.rev15, -automated1, Capella-Gutiérrez et al., 2009), and a maximum-likelihood phylogeny was inferred using IQ-TREE (v2.0.3, -m MFP -B 1000 -alrt 1000 -T AUTO -nm 2000, (Minh et al., 2020). TreeShrink (v1.3.7, -q 0.10, Mai & Mirarab, 2018) was used to remove outlier sequences, and the resulting tree was visualized using iTol (Letunic & Bork, 2024). Clades were manually annotated based on the identity of reference sequences placed within them, and gathering thresholds for the resulting profile HMMs were selected using the same lowest-score/highest-MCC criterion applied throughout this study (see below). New enzyme forms (IV.Aful.a, IV.Aful.b, IV.x) were defined as monophyletic clades. Type II.III sequences, which were not represented among the initial reference sequences, were instead identified via a BLAST search against a single reference sequence obtained from (Banda et al., 2020) and the clade containing the resulting best hit was then annotated as Type II.III. Additionally, Type III sequences were split into three subclades (III.a, III.b, and III.c).

Gathering thresholds for the RuBisCO profile HMMs were determined using stratified 8-fold cross-validation, implemented in Julia (MLDataUtils). For each fold, training sequences were split by subtype, aligned using MAFFT (v7.453, mafft --auto Katoh & Standley, 2013)), and a profile HMM was built from each alignment using HMMER (v3.3.1, hmmbuild, Eddy, 2011). Each initial HMM was searched (v3.3.1, hmmsearch, no threshold, Eddy, 2011) against the concatenated training set, and for each threshold in a candidate range (in increments of 5 bits, spanning the observed sequence score range), true and false positive and negative classifications were tallied and used to calculate the Matthews correlation coefficient (MCC). Then, the lowest threshold achieving the maximum MCC was selected. For subtypes with multiple sub-forms (I, III, IV), sub-form labels were merged to their parent type for this evaluation. This threshold was written into the HMM as its gathering (GA) cutoff and used to evaluate held-out performance on the corresponding validation fold (Figure S2). To derive the final gathering threshold used in production HMMs, the same threshold-selection procedure was applied once to the complete reference sequence set (training and validation sequences combined).

### Profile HMM construction for the remaining nine pathway variants

For the nine remaining pathway variants, reference sequences for diagnostic marker genes were identified from published literature, and orthologous sequences were collected from UniProt using shared PFAMs and KO assignments to capture the full diversity of our genes of interest (true positives), but also retrieve genes that share high sequence similarity and/or functional domains but lack the target catalytic function (false positives). Initial sequence sets were aligned using MAFFT (v7.453, mafft-fftns –adjustdirection --anysymbol, Katoh & Standley, 2013) and maximum-likelihood phylogenies were then inferred from these alignments using IQ-TREE (v2, -m LG+I+G, Minh et al., 2020), with branch support assessed via 1,000 ultrafast bootstrap replicates and 1,000 SH-aLRT replicates. Monophyletic TP clades were defined by inspection of the resulting phylogeny, informed by the known identity of reference sequences placed within each clade. For each TP clade, corresponding sequences were aligned with MAFFT (v7.453, mafft-fftns --adjustdirection, Katoh & Standley, 2013), and separate profile HMMs were built for each TP and FP clade using HMMER (v3.3.1, hmmbuild, Eddy, 2011). For each TP profile HMM, a gathering (GA) threshold was calculated using k-fold cross-validation, with the number of folds limited by the number of available TP sequences maximum ten folds, Table S1). Fold assignment was performed using scikit-learn’s KFold (shuffle=True, random_state=1, VaroquauxGaël & DuchesnayÉdouard, 2011). In each training fold, the sequences used to build that clade’s HMM were treated as TP, and all other related sequences from the same tree, including sequences from other TP variants and designated FP sequences, were combined and treated as FP. The corresponding profile HMM was searched against the training set without a score threshold (HMMer v3.3.1, hmmsearch, Eddy, 2011), and for each fold we identified the lowest bit-score threshold that maximized the Matthews correlation coefficient (MCC), calculated across a scan of candidate thresholds in increments of 5 bits spanning the observed score range. This threshold was then applied to the corresponding held-out test fold (HMMer v3.3.1, hmmsearch, --cut_ga Eddy, 2011) to evaluate held-out performance. The final GA threshold for each clade was calculated as the average threshold across all folds in which the validation MCC exceeded 0.8 (Khedkar et al., 2022), and was written directly into the profile HMM file as its GA cutoff. FP profile HMMs were built as a second, separate set of models from the FP reference clades, using the same alignment and HMM-building procedure (MAFFT v7.453, mafft-fftns --adjustdirection; HMMER v3.3.1, hmmbuild, Eddy, 2011; Katoh & Standley, 2013). Rather than threshold these independently, each FP HMM was assigned the lowest gathering threshold obtained across all TP HMMs, ensuring conservative, permissive detection during downstream screening. To broaden the phylogenetic diversity captured by each TP profile HMM, we screened proGenomes3 (Fullam et al., 2023) with all thresholded TP and FP HMMs, retaining hits that passed their respective clade-specific GA threshold; where a sequence was detected by more than one HMM, only the annotation with the lowest e-value was retained. Identical hit sequences were clustered using MMseqs2 easy-linclust (minimum sequence identity 1.0, coverage 1.0, --cov-mode 0, Steinegger & Söding, 2017), and cluster representatives were aligned to the corresponding reference alignment using MAFFT (v7.453, mafft-fftns --add --keeplength --adjustdirection --anysymbol, Katoh & Standley, 2013). Representative sequences were then phylogenetically placed onto the corresponding reference tree using EPA-ng (v0.3.8, --model LG, Barbera et al., 2019). For each placed sequence, the clade was assigned based on the highest-likelihood-weight-ratio placement, using a custom R script (ape, treeio, tidytree) that propagated reference clade labels through the tree in post-order traversal, assigning “discarded” to internal nodes with conflicting descendant clade identities. This placement-based clade assignment was compared against each sequence’s original HMM-based annotation; only sequences for which the two agreed were retained as confirmed TP hits. Cluster members were subsequently assigned the same classification as their representative sequence. Confirmed proGenomes3 hits were subsequently integrated into the corresponding TP profile HMMs only, and the full k-fold cross-validation procedure described above was repeated on this expanded TP sequence set to re-derive final gathering thresholds (Figure S2).

All final HMMs, reference alignments, and phylogenetic trees generated in this study are provided in the Supp. Data, offering a resource for annotating autotrophic carbon fixation pathways in future genomic and metagenomic datasets.

### Marker gene screening of genomic and metagenomic databases

The resulting, refined HMMs were used to screen predicted genes from SPIRE (Schmidt et al., 2024), proGenomes3 (Fullam et al., 2023), and the MAG catalogue from the Blanes Bay Microbial Observatory time series (Latorre et al., 2025) using HMMER (v3.3.1, hmmsearch --cut_ga, Eddy, 2011), applying the best-hit criterion described above. Putative hits were clustered and confirmed by placement in the reference tree, as described above. For SPIRE specifically, only genes annotated as complete by the database’s precomputed Prodigal gene calls were retained for downstream analysis (Hyatt et al., 2010). Retained hits were classified according to their source into four categories: reference genomes (proGenomes3), low-quality (LQ) MAGs, medium- or high-quality (MQ/HQ) MAGs, and unbinned contigs, following the CheckM2-based quality classifications provided by SPIRE (Chklovski et al., 2023). For pathways represented by more than one marker gene, a genome or MAG was only counted as a positive detection if the required combination of marker genes was present, and, where applicable, if genes indicating alternative or competing pathway variants were absent, according to the co-occurrence and exclusion rules defined for each pathway (Table S1).

### Rarefaction analysis of marker gene diversity

To assess whether the phylogenetic diversity captured across all four detection categories exceeded that captured within genomes and MAGs alone, marker gene sequences were clustered at 95% identity and 80% coverage (--cov-mode 0) using MMseqs2 easy-linclust, to define species-level gene clusters (Steinegger & Söding, 2017). Clustering was performed separately for two sequence sets: the full set of hits across all four detection categories, and hits restricted to reference genomes (proGenomes3) and medium- and high-quality MAGs only. Rarefaction curves were then generated for each set by plotting the cumulative number of clusters recovered against increasing numbers of sampled sequences.

### Phylogenetic, habitat, and physiological characterization of MAGs carrying autotrophic pathways

Taxonomic classifications for reference genomes and medium- and high-quality MAGs were obtained from GTDB v220 (Parks et al., 2022), as provided through SPIRE and proGenomes3. Habitat annotations for samples in which pathways were detected were obtained from (Kim et al., 2026) and further collapsed to broader habitat categories (Table S2). To visualize this distribution, samples were projected onto the pre-existing taxonomic UMAP embedding of (Kim et al., 2026), and coloured by collapsed habitat category and by the presence of each pathway. Habitat enrichment for each pathway was assessed by comparing, for each pathway and habitat cluster, the number of MAGs assigned to that pathway within the habitat against all MAGs not carrying the respective pathway, using two-sided Fisher’s exact tests; resulting p-values were adjusted for multiple testing across all pathway-habitat combinations using the Benjamini-Hochberg (BH) procedure, with significance defined as adjusted p < 0.05. Functional enrichment was assessed using the same Fisher’s exact test procedure, applied to a selected set of metaTraits annotations (Podlesny et al., 2026) of SPIRE MAGs. Oxygen tolerance (obligate aerobic, obligate anaerobic) was annotated using Traitar predictions (Weimann et al., 2016), and all other functional traits were annotated using MICROPHERRET (Bizzotto et al., 2024). Genomes for which Traitar predicted both “obligate aerobic” and “obligate anaerobic” as true were classified as having contradictory oxygen tolerance annotations and excluded from oxygen-status analyses.

### Environmental metadata for marine Campylobacterota MAGs

Environmental metadata for the 196 marine Campylobacterota MAGs, including microntology annotations (Fullam et al., 2026), sampling depth, and dissolved oxygen concentration, were retrieved from the metalog database (Kuhn et al., 2026). Where oxygen data were unavailable, values were instead obtained from World Ocean Atlas climatology (H. E. Garcia et al., 2024), matched to the reported sampling coordinates. Metadata were additionally verified against the corresponding original publications to confirm consistency of reported depth and oxygen concentration (Table S5).

### Functional annotation of Campylobacterota MAGs

General functional annotation of the three Campylobacterota MAGs was performed using eggNOG-mapper v2.1.13 (DIAMOND-based search, eggNOG database v5.0.2, (Buchfink et al., 2021; Cantalapiedra et al., 2021; Huerta-Cepas et al., 2019), and genes involved in nitrogen cycling, sulfur cycling, and hydrogen oxidation were additionally annotated. Respiratory terminal oxidase composition, including the distinction between cbb3- and caa3-type cytochrome oxidases, was determined using METABOLIC v4.0 (-m-cutoff 0.75, KOfam full database, Zhou et al., 2022). Nitrogen cycling genes were identified using DIAMOND v2.1.4 blastp searches against NCycDB, sulfur cycling genes using DIAMOND blastp searches against SCycDB (both e-value < 1e-5, best hit only, Buchfink et al., 2021; Tu et al., 2019; Yu et al., 2021), and hydrogenase genes using HMMER v3.3.1 searches against HydDB (domain score threshold 30, Eddy, 2011; Søndergaard et al., 2016).

Identification of rTCA cycle oxidoreductases required substantial manual curation beyond standard KEGG orthology assignment, since existing KOs do not reliably discriminate between the Por and Oor five-subunit enzyme complexes or their individual subunits. Individual subunits (A, B, D, E, G) were identified using a combination of KO assignment, protein length, gene name, and associated Pfam domain, since no single field reliably identified a given subunit on its own (Table S6). The alpha subunit (porA/oorA) shared a single KO (K00169) with no distinguishing gene name in the eggNOG output. The beta subunit (porB/oorB) shared a single KO (K00170) and was additionally annotated with the correct subunit by gene name (forB or porB). The delta subunit (porD/oorD) was frequently misannotated with KO K02040, the KO for the unrelated phosphate-binding protein pstS, with which it shares the Pfam domain PBP_like_2; true porD/oorD genes were distinguished from genuine pstS hits by the additional presence of the CO_dh Pfam domain, and were inconsistently assigned the gene name forD. The fifth subunit (porE/oorE) was inconsistently annotated under either K00171 or K00172, and, where a gene name was assigned, as forE; its presence was confirmed in all cases by protein length. The gamma subunit (porG/oorG) was identified exclusively via K00172. To reliably discriminate Por from Oor enzymes, complex identity was established by phylogenetic placement of the alpha subunit. Alpha subunit sequences from the three MAGs and *S. pluma* were aligned to a set of reference sequences using MAFFT (v7.453, mafft-fftns --adjustdirection --anysymbol, (Katoh & Standley, 2013) and used to infer a maximum-likelihood phylogeny in IQ-TREE (-m LG+I+G, 1,000 ultrafast bootstrap and 1,000 SH-aLRT replicates, Minh et al., 2020) for the main figure (Figure 4B). All ten subunit-encoding genes (five Por, five Oor) were consistently found on the same contig in each genome, organized as two adjacent groups of five separated by a gap; identity of the beta, delta, epsilon, and gamma subunits in each operon was therefore inferred from genomic co-location with the alpha subunit gene already classified by phylogenetic placement. Within this reference set, we additionally noted that the gene names forB2 and forD were assigned by eggNOG-mapper exclusively to Oor subunits, corroborating the alpha-subunit-based classification, though we did not verify whether this naming pattern held more broadly and did not rely on it for classification. Genes encoding bacterial globins and hemerythrins co-locating with oxygen-tolerant oxidoreductases were identified through manual screening of eggNOG-mapper annotations on the contigs carrying the five-subunit por and oor genes.

Building on these individual gene-level annotations, we summarized presence and absence across the same set of physiological marker genes examined by (Molari et al., 2023) into high-level functional-category calls per MAG (Note S2, Table S7, (Molari et al., 2023).

To compare the MAGs from the surface ocean to the previously described *S. pluma* and to assess whether the features observed in the three MAGs were typical or unusual within their respective genera, we compared annotations from the two *S. plum*a MAGs identified proGenomes3 reference genomes (Fullam et al., 2023) sharing the same GTDB r220 (Parks et al., 2022) genus annotations as each MAG. *S. pluma* and progenimes3 references were annotated using eggNOG-mapper v2.1.13 and METABOLIC v4.0 as described above. The porA/oorA phylogeny was re-inferred including sequences annotated in these reference genomes (Figure S1), as described above, to determine whether each carried the five-subunit or canonical four-subunit form of Por/Oor. For each reference genome, we further recorded the presence of a caa3- or cbb3-type cytochrome oxidase, the presence of dissimilatory nitrate reduction genes (napA, napB), whether the por and oor operons occurred on the same contig, and, where co-located, the number of genes separating the two operons (Table S8).

### Metagenomic and metatranscriptomic read mapping of TARA Oceans against Campylobacterota MAGs

Metagenomic reads from the TARA Oceans dataset were mapped against the three MAGs using CoverM (CoverM genome, bwa-mem mapper, minimum read identity 95%, covered_fraction and trimmed_mean methods, Aroney et al., 2025), and a MAG was considered detected in a sample when at least 50% of its genome length was covered. Metatranscriptomic reads were quality- and adapter-trimmed using BBDuk (BBMap v38.26; ktrim=r k=23 mink=11 hdist=1 qtrim=rl trimq=10 minlen=36, Bushnell B.: BBMap. sourceforge.net/projects/bbmap) and mapped against each MAG’s predicted gene set using kallisto (v0.51.1, kallisto quant, Bray et al., 2016). A MAG was considered transcriptionally active in a sample when at least 10 annotated genes had non-zero mapped reads. To additionally assess coverage of individual rTCA pathway genes (Supp. Data) metagenomic reads were also mapped at the gene level using BBMap (minid=0.95, ambiguous reads assigned randomly), with per-gene coverage and depth summarized using pileup.sh (Bushnell B.: BBMap. sourceforge.net/projects/bbmap/).

To compare the environmental conditions (depth, nitrate, and dissolved oxygen concentration) under which each MAG was detected against the full set of screened samples, we used Kruskal-Wallis tests followed by Dunn’s post hoc test with Benjamini-Hochberg correction for multiple comparisons (Table S9). To test whether transcriptionally active samples differed from all coverage-confirmed samples for the same variables, we used two-sided Wilcoxon rank-sum tests per MAG and variable, with Benjamini-Hochberg correction applied across all tests.

## Supporting information

Supplementary Notes, Figures and Table headers

## Availability of additional data and scripts

All materials and data generated during this study are shared under https://doi.org/10.5281/zenodo.23211329.

## Funding and support

Funded by the European Union under the Horizon Europe Programme, Grant Agreement No. 101082304 (BlueRemediomics). Views and opinions expressed are however those of the author(s) only and do not necessarily reflect those of the European Union or the granting authority, the Research Executive Agency (REA). Neither the European Union nor the granting authority can be held responsible for them.

This work was supported by the EMBL IT Services HPC resources (DOI 10.5281/zenodo.12785829).

We thank the Blanes Bay Microbial Observatory team for sample collection, the projects supporting the initiative (https://bbmo.icm.csic.es), and Francisco Latorre and Lidia Montiel for their contributions to generating the Blanes Bay MAGs.

## Author contributions

AM and PB designed the study.

AM wrote the manuscript with input from all authors.

AM and SW built and trained the profileHMMs and AM designed and ran the final search pipeline with input from AF, JHC, and DE.

AM performed data analysis with input from JS and RL.

MK and PB supervised the study.

## Competing interests

The authors declare no competing interests.

## Declaration of generative AI and AI-assisted technologies in the writing process

During the preparation of this work, the authors used Claude (Anthropic) in order to assist with coding as well as language editing and improving clarity. After using this tool, the authors reviewed and edited the content as needed and take full responsibility for the content of the published article.

## Notes

### Competing Interest Statement

The authors have declared no competing interest.

### Summary of Updates

Added a link to supplementary data and additional materials, update of the results

