## Supplementary Notes, Figures and Table headers for "A global genomic survey of prokaryotic carbon fixation reveals an oxygen-tolerant rTCA cycle in the surface ocean"

† Deceased

* Corresponding authors

*Content*

Contains or links to:

Supplementary Note 1-2

Supplementary Figures 1-12

Supplementary Tables 1-9

*Notes*

**Note S1: Oxygen-associated trait enrichment across rTCA-carrying phyla**

To characterize oxygen-related physiology at phylum resolution across the three rTCA cycle variants, we examined the prevalence of aerotolerance, obligate aerobicity, and obligate anaerobicity within each pathway's dominant phyla (for phyla with at least ten MAGs, Table S4). Within rTCA1b, Aquificota were uniformly obligate anaerobic (100% of trait-annotated genomes), matching this lineage's classical association with anoxic and microoxic thermal environments (Hügler & Sievert, 2011). Within rTCA1a, Nitrospirota were predominantly both nitrifying (95.2%) and obligate aerobic (95.6%), with obligate anaerobicity being rare (1.3%), matching their established ecology as aerobic nitrifiers (Bayer et al., 2021; Lücker et al., 2010). Within rTCA2, Aquificota and SZUA-79 showed a consistent anaerobic signature (both 100% obligate anaerobic, 0% aerotolerant), while Nitrospirota_A were uniformly obligate aerobic (100%) but, unlike Nitrospirota in rTCA1a, showed no meaningful prevalence of nitrification (0%).

One result, however, does not fit expectations, and we flag it explicitly as a likely limitation of the underlying trait prediction rather than a genuine biological signal. Within rTCA1a, Bacteroidota showed a markedly different oxygen-tolerance profile from the pathway's other phyla: 68.1% were aerotolerant, 59.6% obligate aerobic, and 19.1% obligate anaerobic, representing a diverse, internally mixed pattern spanning all three categories, in contrast to the more uniform signatures seen in the other phyla described above. This is inconsistent with the phylogenetic placement of rTCA1a-carrying Bacteroidota, which are exclusively members of Chlorobiaceae (predominantly *Chlorobium*; Supp. Data). Chlorobiaceae are known as green sulfur bacteria with a well-established obligate anaerobic, photolithoautotrophic physiology typically associated with stratified, sulfidic, light-exposed water columns and fundamentally incompatible with sustained aerobic growth (S. L. Garcia et al., 2021). A majority of these genomes nonetheless received "obligate aerobic" calls. Given the strength of the physiological and ecological evidence for obligate anaerobiosis in this lineage, we consider this pattern most likely to reflect a limitation of the genome-based oxygen-tolerance prediction specific to this lineage, plausibly due to sparse or unrepresentative training data for this ecological group, rather than a true departure from expected physiology. We therefore treat the obligate aerobic/anaerobic enrichment result for this phylum with caution and do not report it in the main text.

**Note S2: Summary of the functional annotations of MAG BB, MAG HL, and MAG IO**

**Hydrogen oxidation** All three MAGs encode multiple copies of NiFe-hydrogenases (Table S6), indicating hydrogen could be one electron donor for all three lineages alongside sulfur compounds.

**Sulfur metabolism** All three MAGs encode a complete Sox system, consistent with a capacity for sulfur oxidation. Assimilatory sulfate reduction genes are present only in MAG HL (CysD, CysN, CysH, CysC), while all three MAGs share sir, encoding an alternative assimilatory sulfite reductase; CysI is absent in all three. Genes required for dissimilatory sulfate/sulfite reduction (AprA, AprB, DsrA, DsrB) are entirely absent in all three MAGs, and all three additionally lack sulfur reduction capacity (PsrA/PsrB/PsrC-NrfD entirely absent). Reduced sulfur compounds therefore serve as an electron donor rather than a terminal electron acceptor in these organisms.

**Nitrogen metabolism** MAG BB retains dissimilatory nitrate reduction genes (NapA, NapB present) but lacks NorB, precluding complete denitrification to N2. MAG HL and MAG IO largely lack dissimilatory nitrate reduction genes instead (NapA absent in both; NapB additionally absent in HL), consistent with their oxygen-respiring strategy. Genes for nitrate uptake and assimilation were detected more variably. NarK is absent in MAG BB but present in both MAG HL and MAG IO, while NarB shows the opposite pattern, present in MAG BB and MAG IO but absent in MAG HL. NirA, reducing nitrite to ammonium, is present in all three MAGs. Together, these genes suggest some capacity for nitrate assimilation across all three MAGs, though the specific complement of transport and reduction genes varies.

**Oxygen respiration** MAG HL and MAG IO both encode an additional caa3-type cytochrome oxidase alongside the ancestral cbb3-type oxidase typical of Campylobacterota; MAG BB retains only cbb3-type respiration but additionally encodes a bd-type oxidase, likewise associated with high oxygen affinity. All three MAGs thus supplement or replace the ancestral cbb3 system with a high-oxygen-affinity alternative, despite doing so through different enzymatic routes.

**rTCA cycle** In addition to the complete five-subunit por and oor operons, all three MAGs also encode the pathway specific ATP citrate lyase (aclA, aclB), not only detected by our own HMM pipeline but additionally confirmed through annotations by eggnog and METABOLIC.

*Figures*

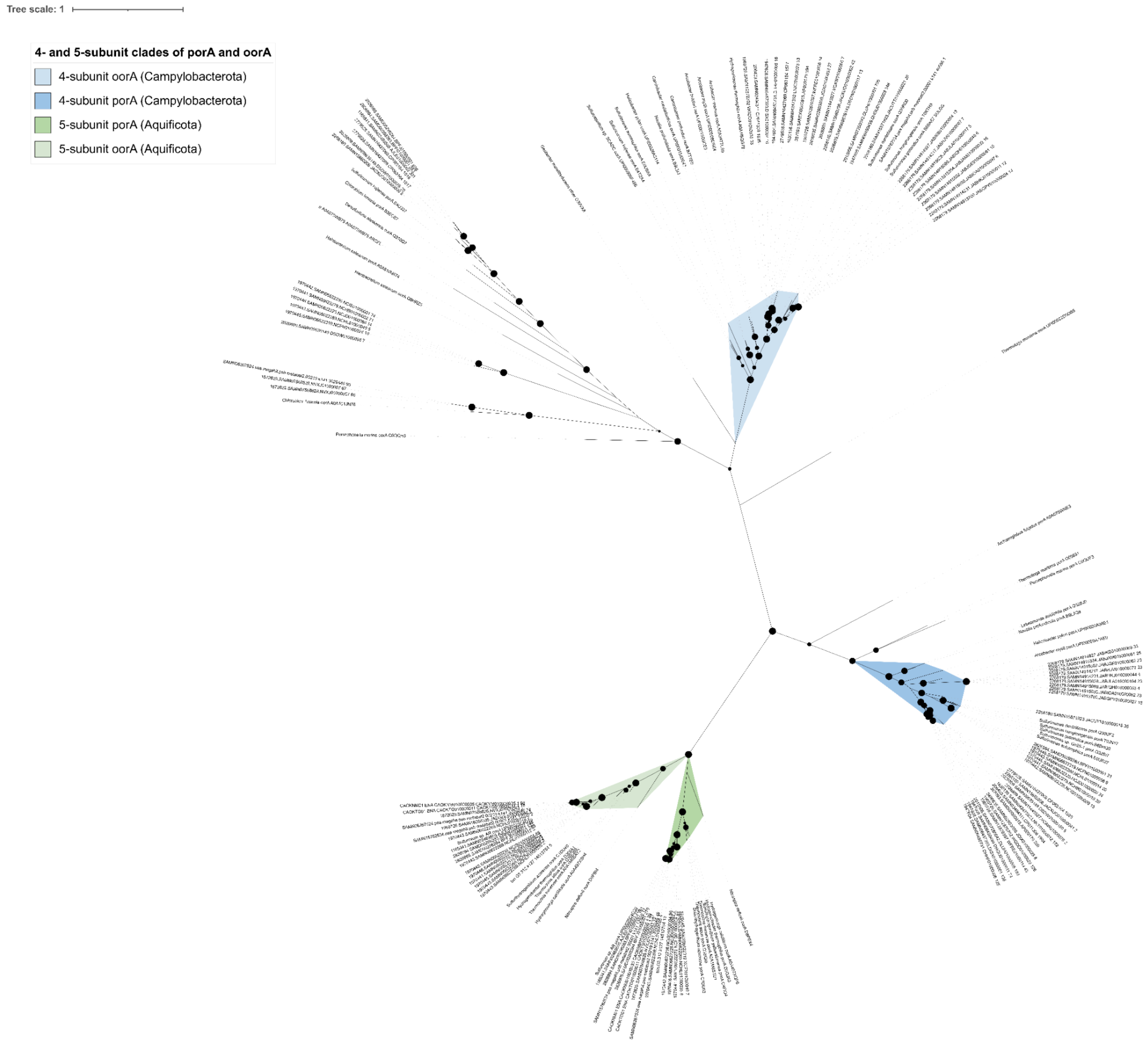


**Figure S1 | Extended phylogeny of the Por/Oor alpha subunits (porA, oorA), including the three MAGs of interest and genus-matched proGenomes3 reference genomes.** This is the full tree underlying the main-text summary shown in Figure 4B, built using the same maximum-likelihood approach on MAFFT-aligned sequences, with four- and five-subunit variants distinguished by phylogenetic placement and confirmation of the complete five-gene operon (Methods). Bootstrap support >= 90 is indicated by black dots. An interactive version of this tree is available at [https://itol.embl.de/tree/101113563236511779459232](https://itol.embl.de/tree/101113563236511779459232).

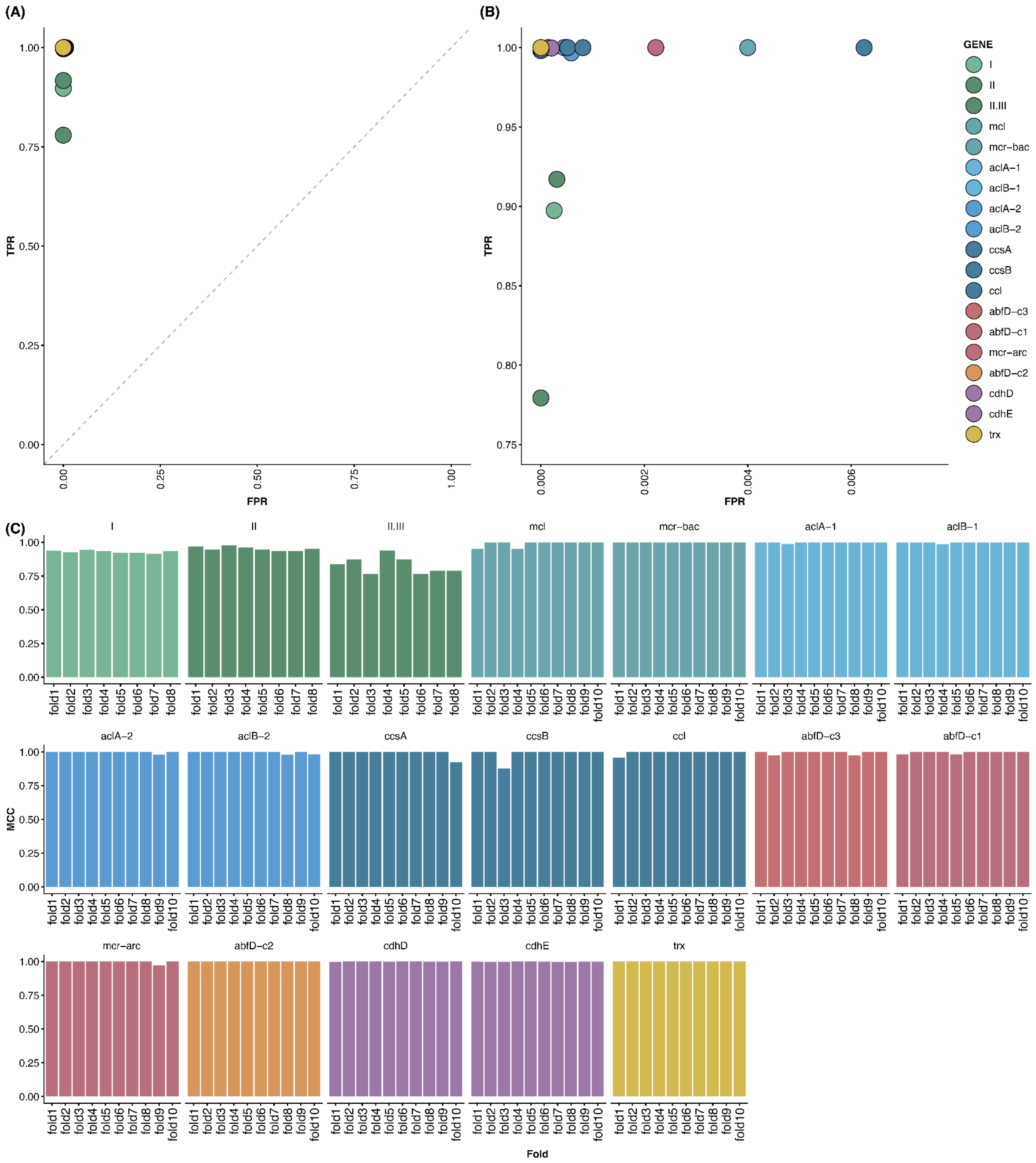


**Figure S2 | Cross-validated classification performance of marker gene profile HMMs.** (A) True positive rate (TPR) against the false positive rate (FPR) for each marker gene's final profile HMM, averaged across cross-validation folds where Matthews correlation coefficient (MCC) was ≥ 0.8. (B) Zoomed view of the high-TPR and low-FPR region (TPR≥ 0.75, FPR ≤ 0.0075) from (A), highlighting classification performance near the operating threshold. (C) MCCs for each marker gene, shown per cross-validation fold.

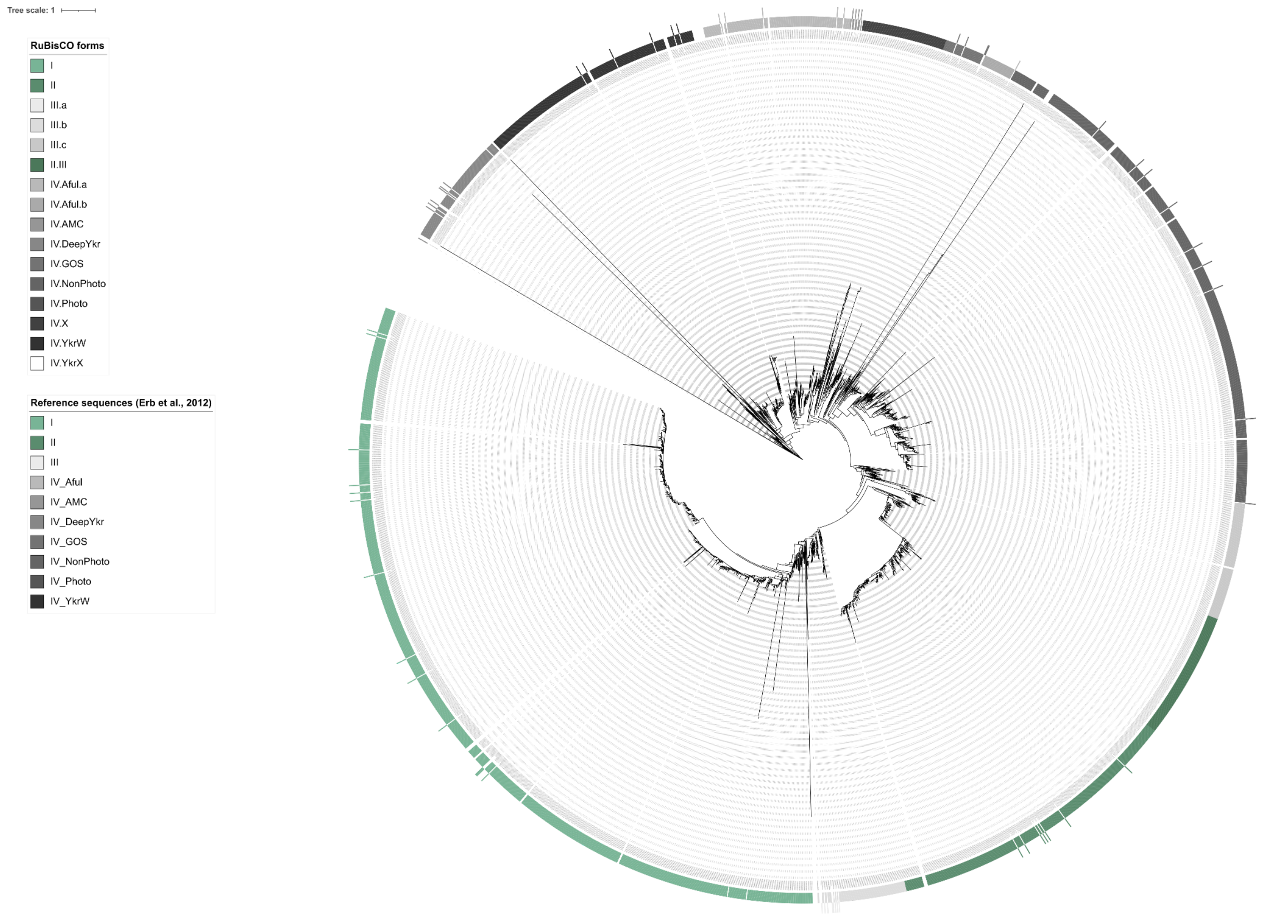


**Figure S3 | A maximum-likelihood phylogenetic tree of RuBisCO large subunit sequences detected in proGenomes3** (Fullam et al., 2023) **and reference sequences derived from** (Erb et al., 2012)**.** RuBisCO forms are indicated by the colored rings (inner ring: proGenomes3 sequences, outer ring: reference sequences). Form I, II, and II.III were used as markers for the CBB cycle while other forms were used as false positive sequences for our search pipeline.

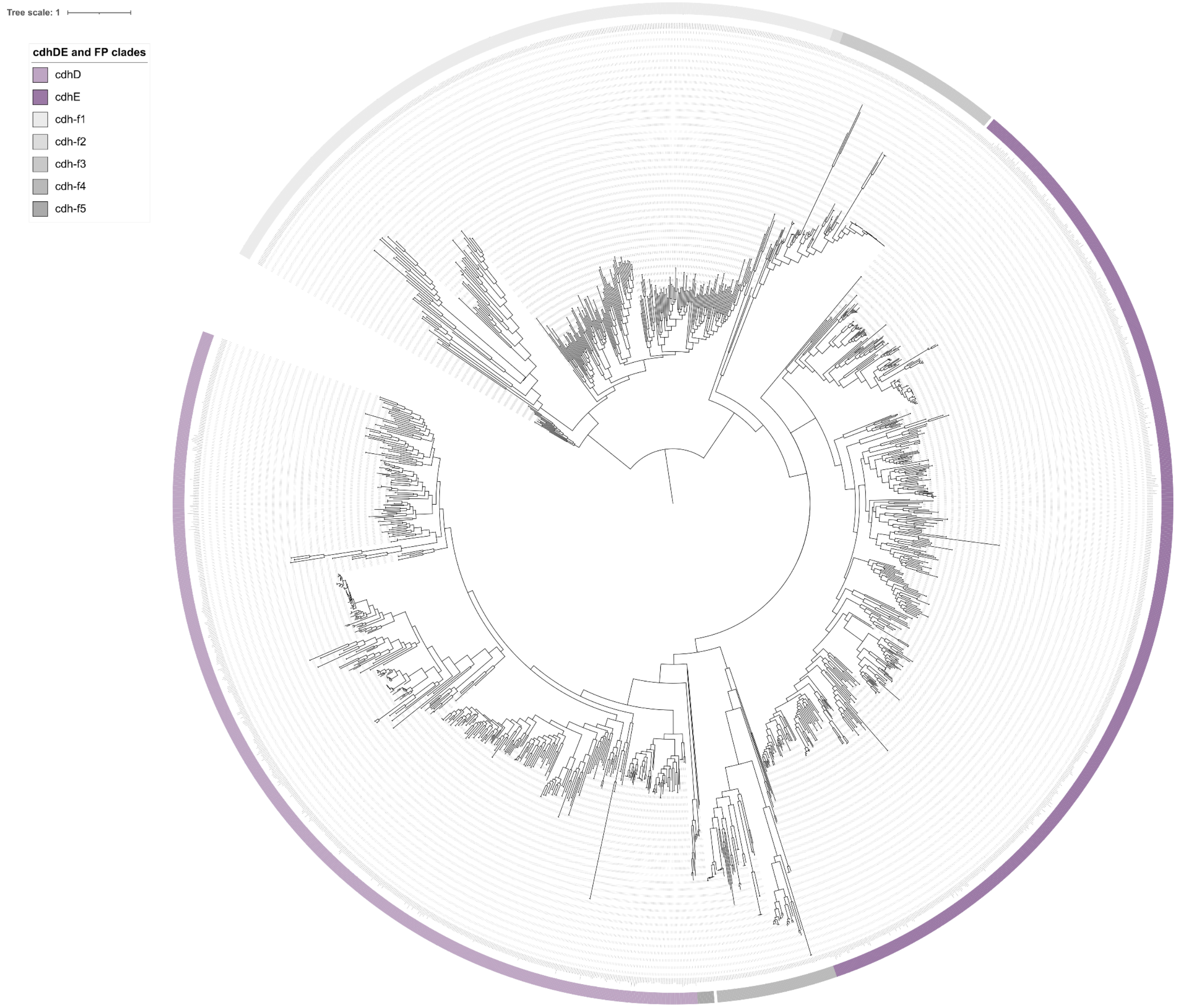


**Figure S4 | A maximum-likelihood phylogenetic tree of cdhD and cdhE genes, as well as sequences sharing either the same KO annotation or PFAM domains (Methods).** Clades are indicated by the colored rings.

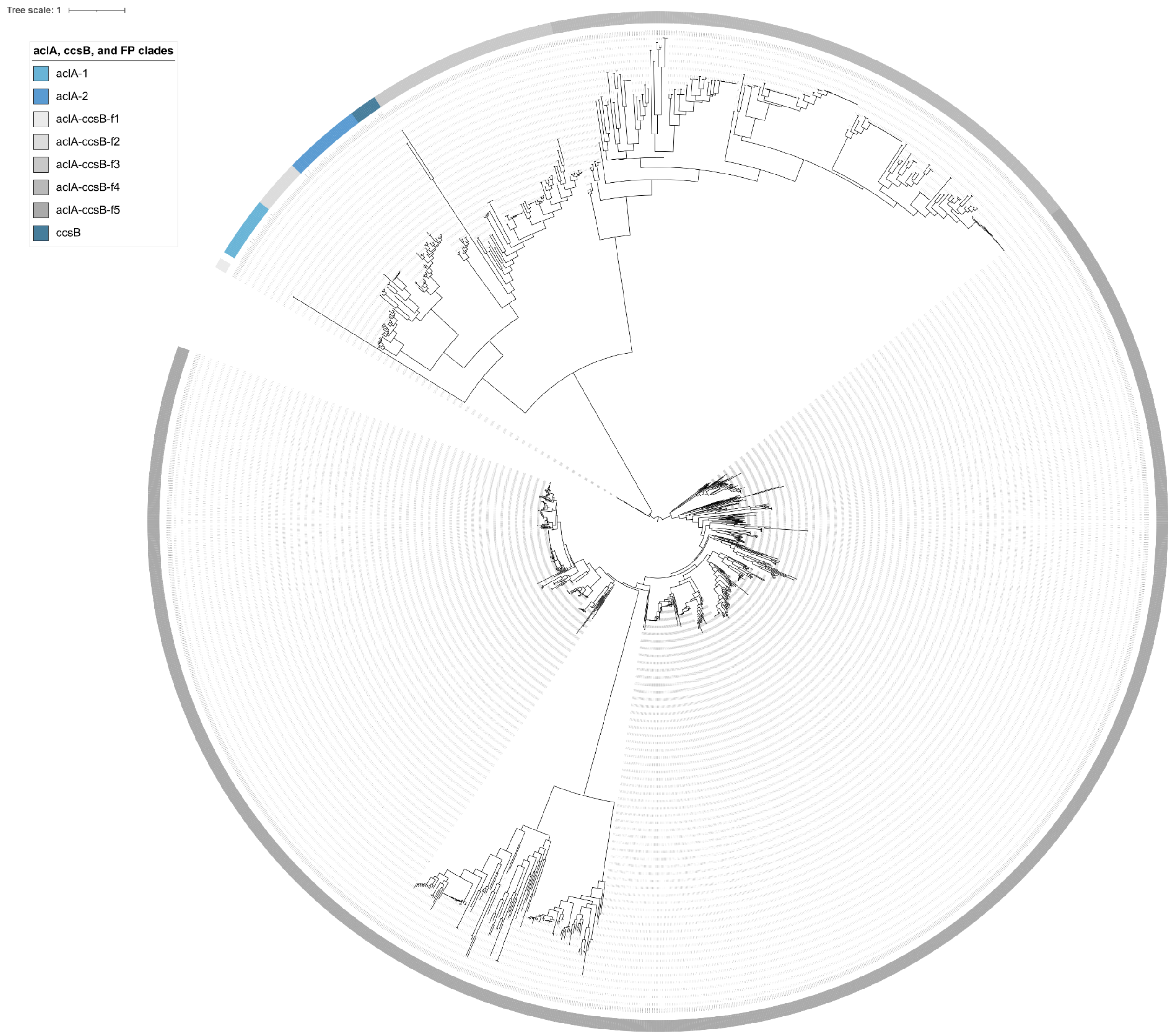


**Figure S5 | A maximum-likelihood phylogenetic tree of aclA and ccsB genes, as well as sequences sharing either the same KO annotation or PFAM domains (Methods).** Clades are indicated by the colored rings.

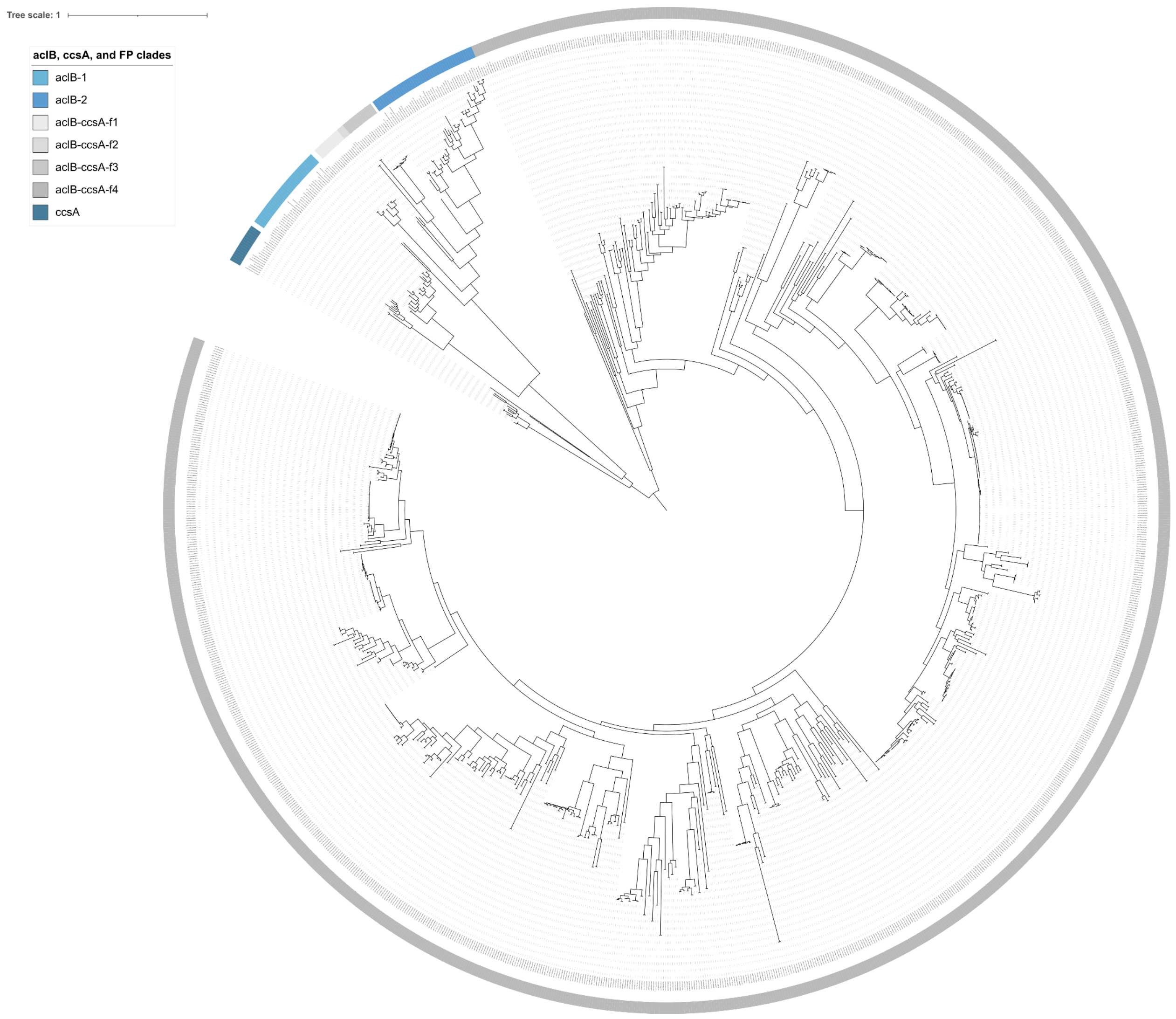


**Figure S6 | A maximum-likelihood phylogenetic tree of aclB and ccsA genes, as well as sequences sharing either the same KO annotation or PFAM domains (Methods).** Clades are indicated by the colored rings.

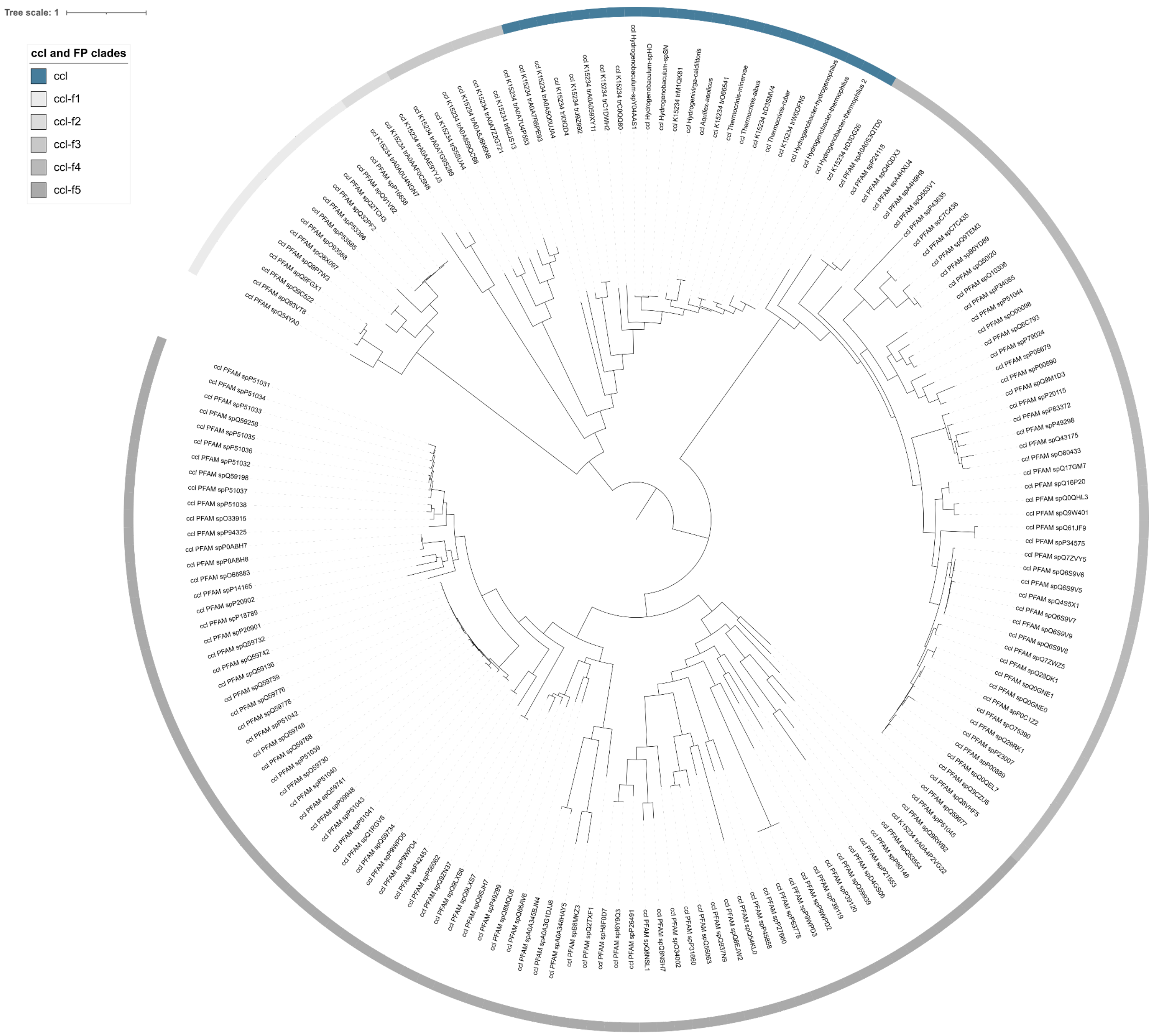


**Figure S7 | A maximum-likelihood phylogenetic tree of ccl genes, as well as sequences sharing either the same KO annotation or PFAM domains (Methods).** Clades are indicated by the colored rings.

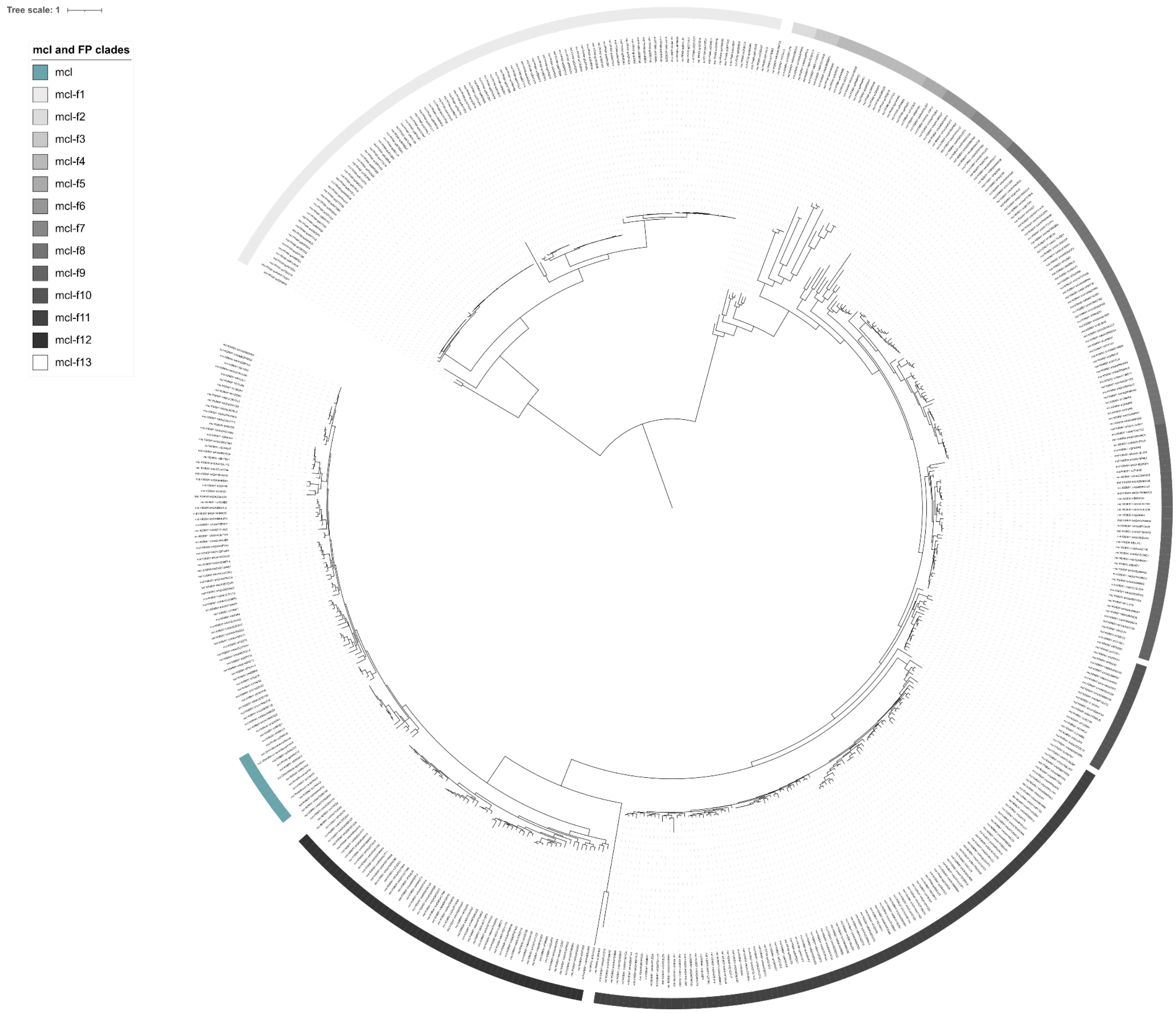


**Figure S8 | A maximum-likelihood phylogenetic tree of mcl genes, as well as sequences sharing either the same KO annotation or PFAM domains (Methods).** Clades are indicated by the colored rings.

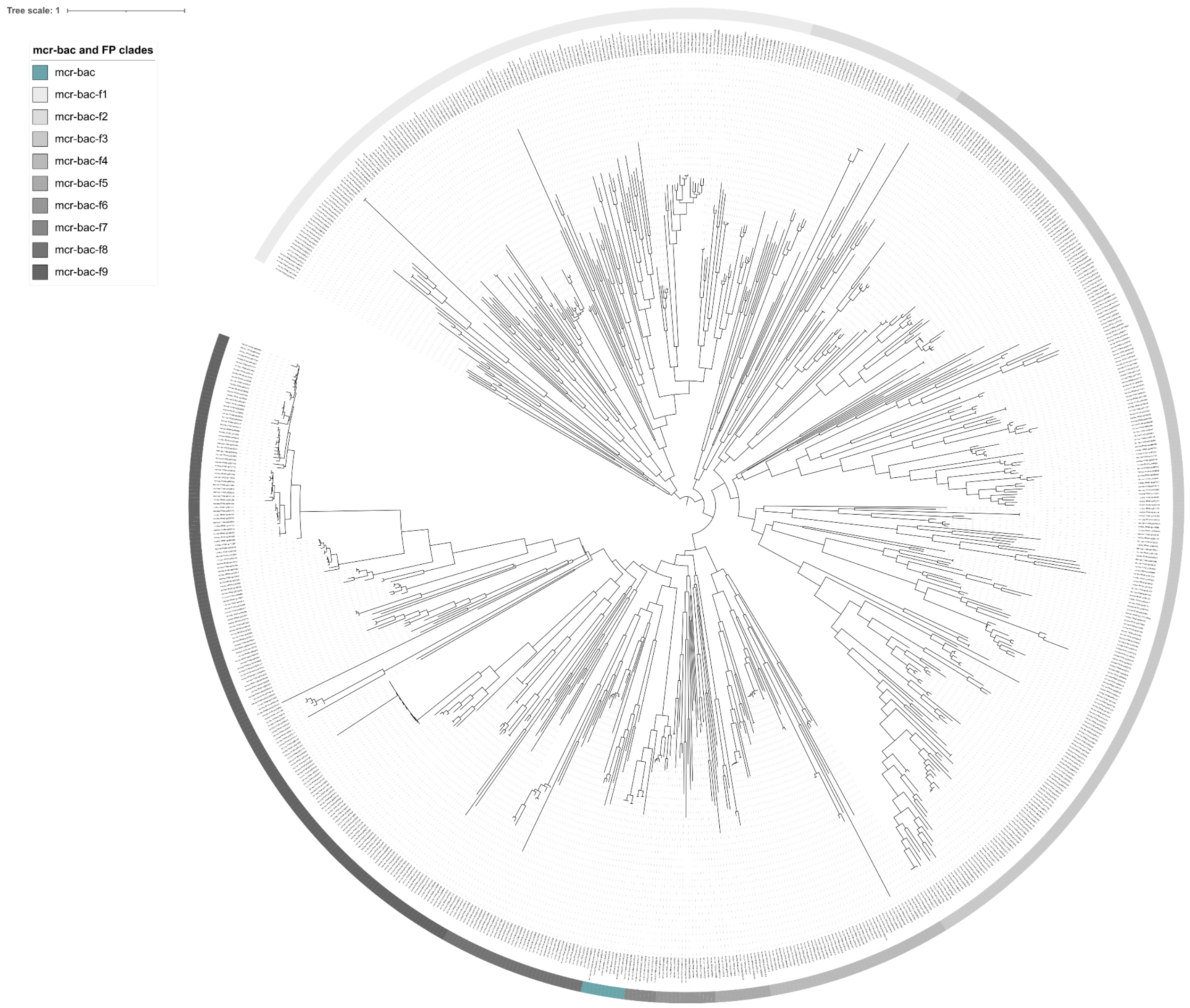


**Figure S9 | A maximum-likelihood phylogenetic tree of mcr-bac genes, as well as sequences sharing either the same KO annotation or PFAM domains (Methods).** Clades are indicated by the colored rings.

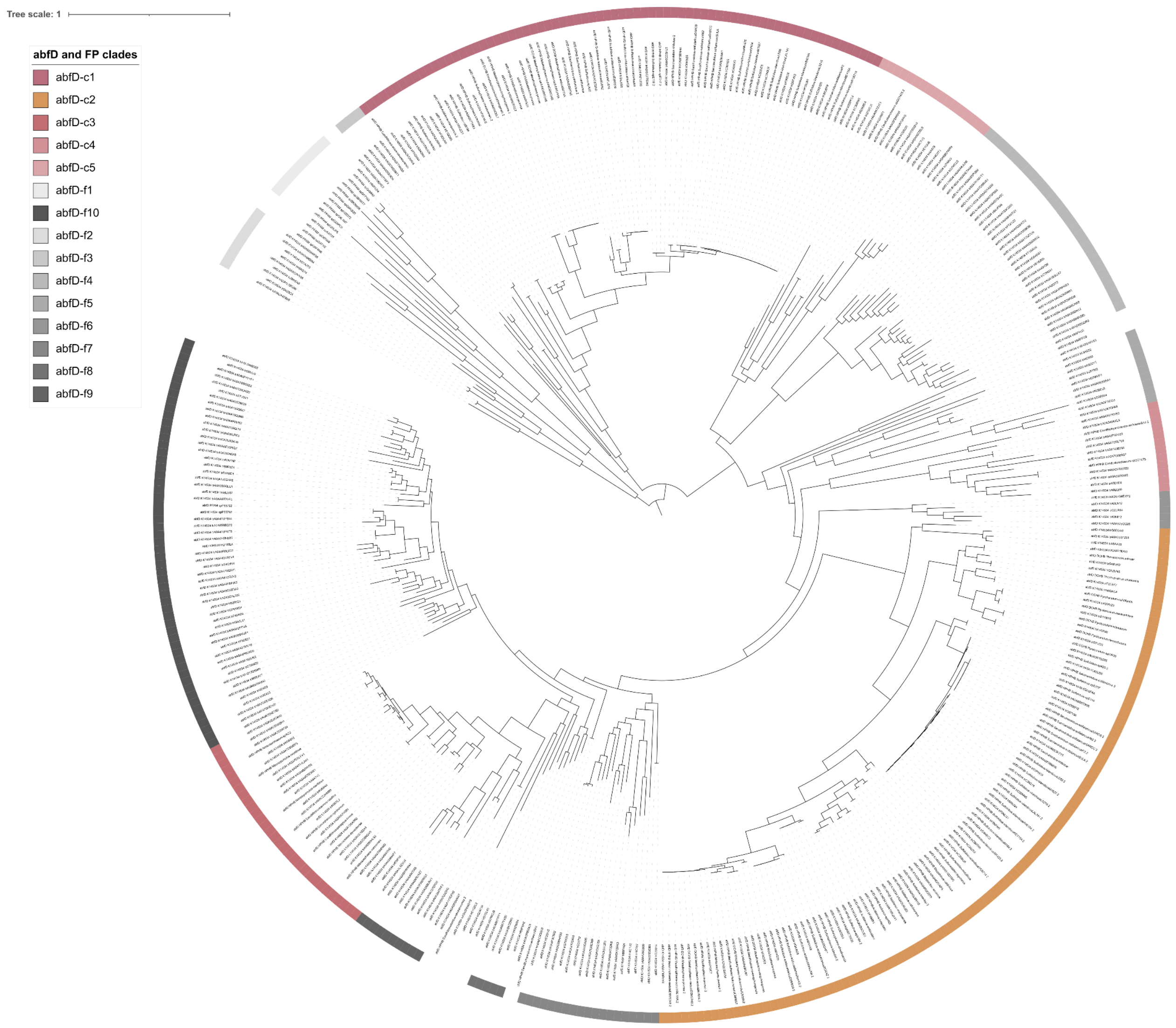


**Figure S10 | A maximum-likelihood phylogenetic tree of abfD genes, as well as sequences sharing either the same KO annotation or PFAM domains (Methods).** Clades are indicated by the colored rings.

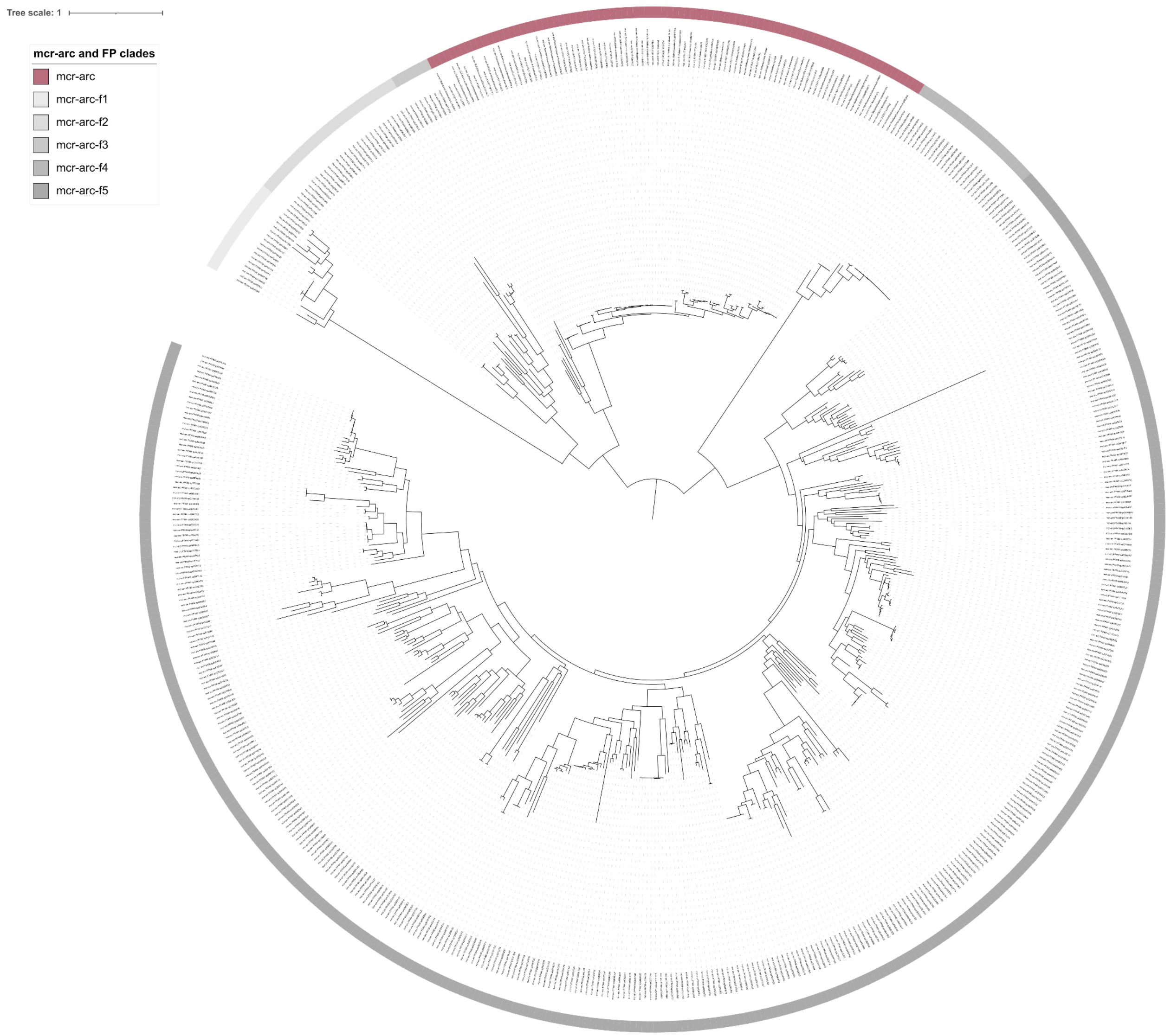


**Figure S11 | A maximum-likelihood phylogenetic tree of mcr-arc genes, as well as sequences sharing either the same KO annotation or PFAM domains (Methods).** Clades are indicated by the colored rings.

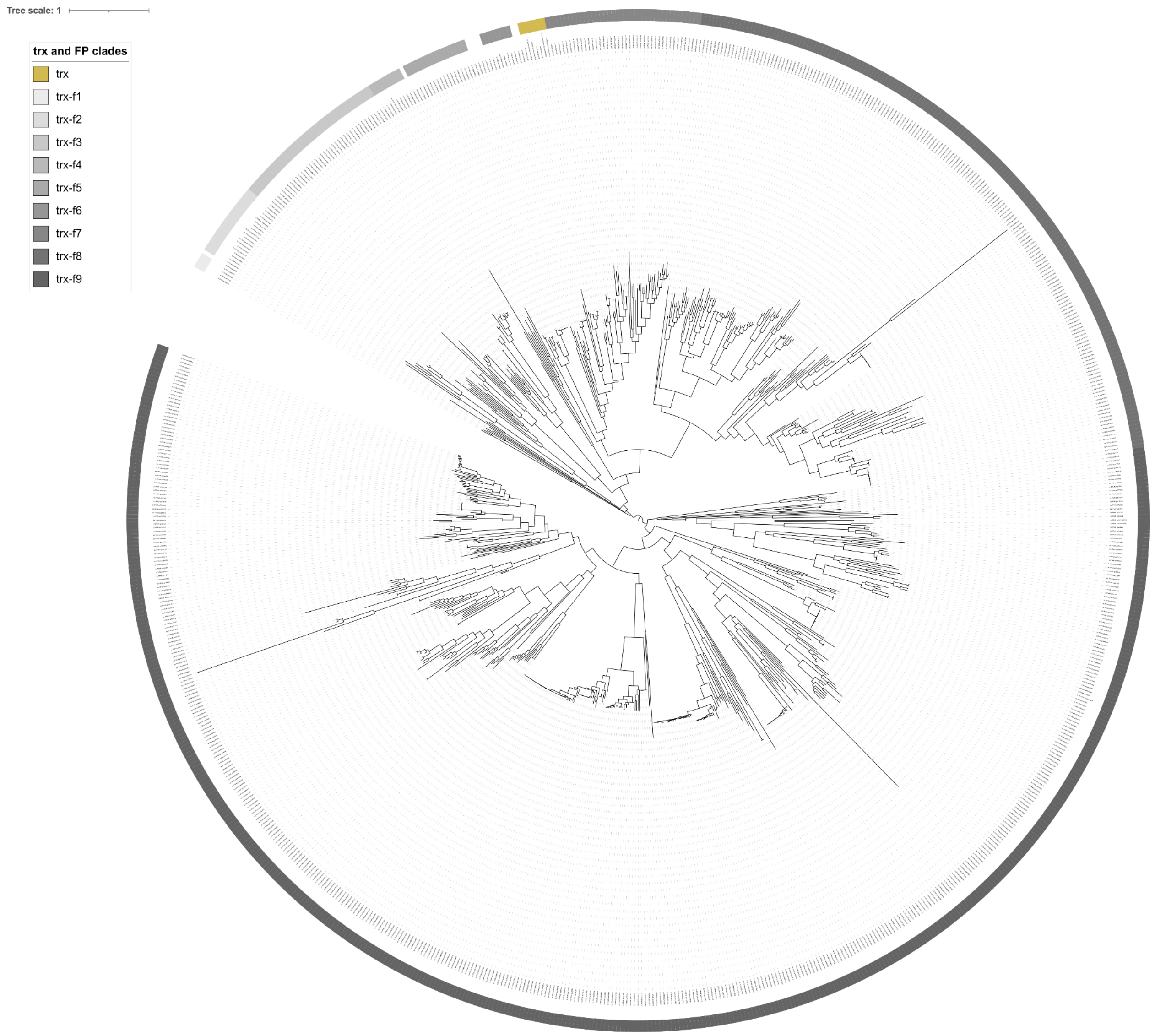


**Figure S12 | A maximum-likelihood phylogenetic tree of trx genes, as well as sequences sharing either the same KO annotation or PFAM domains (Methods).** Clades are indicated by the colored rings.

*Tables*

All supplemental tables are shared via a separate file.

**Table S1 | Carbon fixation pathways, variants, and the marker genes used for their detection.** For each pathway variant, the table lists the marker gene(s) used, the corresponding phylogenetic reference tree (provided as a supplementary figure, with an interactive iTOL version linked), and the criteria required for genome/MAG-level pathway detection (e.g., co-occurrence of multiple marker genes, or presence/absence combinations distinguishing closely related variants). Remarks note pathway-specific caveats identified during marker gene validation, including borderline cases resolved through co-occurrence logic (3-HPB) and the absence of an established marker gene set requiring a custom, phylogenetically restricted approach (rG pathway; Methods).

**Table S2 | Habitat categories defined by** (Kim et al., 2026) **and their combined categories used in this study.** Each fine-grained habitat cluster from the original UMAP-based classification is listed with its original cluster ID and description, alongside the broader combined category into which it was collapsed for the enrichment analyses presented here (Figure 2E; Methods).

**Table S3 | Habitat and functional enrichment for each carbon fixation pathway.** For each pathway, all tested habitat categories (Table S2) and physiological traits are shown alongside Odds ratios, raw and Benjamini-Hochberg-adjusted p-values, and significance calls (two-sided Fisher's exact test comparing each pathway against all other pathways; significance defined as adjusted $p < 0.05$). This table reports all tested pathway–habitat and pathway–trait combinations, including those not reaching significance, and underlies the enrichment results shown in Figures 2E and 2F.

**Table S4 | Habitat and functional enrichment for the three most common phyla within each carbon fixation pathway.** For each pathway, the three phyla with the most MAGs are shown alongside the number of MAGs in that phylum, and all tested habitat categories (Table S2) and physiological traits, with Odds ratios, raw and Benjamini-Hochberg-adjusted p-values, and significance calls (two-sided Fisher's exact test comparing each pathway against all other pathways; significance defined as adjusted $p < 0.05$). This table reports all tested phylum–habitat and phylum–trait combinations within each pathway, including those not reaching significance, and underlies the phylum-resolved results discussed in the main text. Odds ratios reported as 'Inf' reflect a zero-count cell in the underlying contingency table (i.e., the trait or habitat was detected in only one of the compared groups) and indicate complete separation rather than a quantifiable effect size.

**Table S5: Habitat description, depths, and oxygen concentrations at sampling sites where marine Campylobacterota were detected.** Italic entries indicate WOA modeled values that do not match the microntology or the sample type based on the study description (e.g., methane seeps and estuaries). Samples which contained the MAGs further analyzed are highlighted in blue and bold.

**Table S6: Gene-level annotation evidence for the Por and Oor five-subunit oxidoreductase complexes in the three Campylobacterota MAGs of interest.** For each subunit-encoding gene, the table lists the KO assignment(s), associated Pfam domain(s), and protein length used to identify subunit identity, alongside the phylogenetic classification of the alpha subunit (porA/oorA) and the final complex/subunit annotation assigned by combining KO, Pfam, length, gene name, and genomic co-location, as described in the Methods. Genes encoding co-located bacterial globins and hemerythrins are also included. This table underlies the manual curation approach described in the main text and Methods for discriminating Por from Oor complexes and their individual subunits.

**Table S7 | Presence of marker genes for key metabolic functions in the three Campylobacterota MAGs analyzed in this study.** Genes are grouped by functional category (hydrogen oxidation, sulfur cycling, nitrogen cycling, oxygen respiration, and the rTCA cycle), gene copy number shown for each MAG. This table underlies the functional annotation summary presented in the main text and in Note S2.

**Table S8 | Comparative genomic summary of oxygen-tolerance-associated features across the *S. pluma* reference MAGs, the three MAGs of interest from this study, and proGenomes3 reference genomes matching the genus of the three MAGs of interest (based on GTDB r220 annotations).** For each genome, the table reports presence of the five-subunit Por and Oor variants, whether porA and oorA are located on the same contig and, if so, the number of genes separating them, and presence of dissimilatory nitrate reductase (napAB) and the cbb3- and caa3-type cytochrome c oxidases. Yes/no values are colour-coded (green: yes, red: no), with the three MAGs of interest shown in darker shades and reference genomes in lighter shades for visual distinction.

**Table S9 | Spearman correlations between genome coverage metrics and environmental variables across TARA Oceans samples.** For MAG BB and MAG HL, correlations between percent genome covered and mean sequencing coverage against six environmental parameters (depth, nitrate, chlorophyll, oxygen, salinity, and temperature) are shown, along with sample size, Spearman's rho, raw p-value, and Benjamini-Hochberg-adjusted p-value. MAG IO was excluded from this analysis due to its low number of detected samples (n=2), insufficient for meaningful correlation testing.

*Data*

All Supp. Data is available under https://doi.org/10.5281/zenodo.23211329.
